# Female GluA3-KO mice show early onset hearing loss and afferent swellings in ambient sound levels

**DOI:** 10.1101/2024.02.21.581467

**Authors:** Indra Pal, Atri Bhattacharyya, Babak V-Ghaffari, Essence D. Williams, Maolei Xiao, Mark A. Rutherford, María Eulalia Rubio

## Abstract

AMPA-type glutamate receptors (AMPAR) mediate excitatory cochlear transmission. However, the unique roles of AMPAR subunits are unresolved. Lack of subunit GluA3 (*Gria3^KO^*) in male mice reduced cochlear output by 8-weeks of age. Since *Gria3* is X-linked and considering sex differences in hearing vulnerability, we hypothesized accelerated presbycusis in *Gria3^KO^* females. Here, auditory brainstem responses (ABR) were similar in 3-week-old female *Gria3^WT^*and *Gria3^KO^* mice. However, when raised in ambient sound, ABR thresholds were elevated and wave-1 amplitudes were diminished at 5-weeks and older in *Gria3^KO^*. In contrast, these metrics were similar between genotypes when raised in quiet. Paired synapses were similar in number, but lone ribbons and ribbonless synapses were increased in female *Gria3^KO^*mice in ambient sound compared to *Gria3^WT^* or to either genotype raised in quiet. Synaptic GluA4:GluA2 ratios increased relative to *Gria3^WT^*, particularly in ambient sound, suggesting an activity-dependent increase in calcium-permeable AMPARs in *Gria3^KO^*. Swollen afferent terminals were observed by 5-weeks only in *Gria3^KO^* females reared in ambient sound. We propose that lack of GluA3 induces sex-dependent vulnerability to AMPAR-mediated excitotoxicity.

## Introduction

The α-amino-3-hydroxy-5-methyl-4-isoxazolepropionic acid receptors (AMPARs) are tetrameric glutamate-gated ion channels with pore-forming subunits GluA1-4 encoded by a family of genes, *Gria1-4*. Each subunit contributes to ionic permeability, pharmacological sensitivity, intracellular trafficking, synapse development, and activity-dependent plasticity of the channel. GluA1 and GluA2 are the most abundant subunits, relatively ubiquitous in the central nervous system, and prominently determining AMPAR plasticity and function, respectively (Shanks et al., 2012; Bassani et al., 2013; Purkey and Dell’Acqua, 2020). In particular, the absence of GluA2 from the AMPAR tetramer increases the unitary current and makes it permeable to Ca^2+^ in addition to Na^+^ ions. In the adult hippocampus and cortex, the canonical Ca^2+^-permeable AMPAR (CP-AMPAR) involved in synaptic plasticity is the GluA1 homomeric channel. GluA3 and GluA4 have more unique expression patterns, tending to be found at synapses with relatively fast kinetics, such as those in the cochlea and auditory brainstem (Wang et al., 1998; Rubio et al., 2017; Rutherford et al., 2023). Interestingly, the mature cochlea and auditory brainstem do not express GluA1, but they do express CP-AMPARs comprised of GluA3 and GluA4 subunits.

In male mice, the pore-forming subunit GluA3 of the AMPAR is essential for the development and maintenance of the cochlear afferent ‘ribbon’ synapses that underlie hearing function. Wild-type and GluA3^KO^ mice raised in ambient sound levels had similar hearing sensitivities at one month of age, although ribbon synapses already exhibited ultrastructural and molecular alterations. These alterations include defective position-dependent maturation of pre- and post-synaptic specializations along an axis of structural and functional differentiation, as well as an imbalance of post-synaptic GluA subunit expression – an increase in GluA4 and decrease in GluA2 – suggesting a shift toward greater permeability to Ca^2+^ (Rutherford et al., 2023). At 2 months, the cochleae of male GluA3^KO^ mice were less sensitive to suprathreshold sounds (lower wave-1 amplitude of the Auditory Brainstem Response, ABR), and by 3 months of age, they were less excitable by low-level sounds (elevated ABR threshold; García-Hernández et al., 2017). These results showed that synaptic anatomical defects preceded synapse dysfunction leading to hearing loss in male mice lacking GluA3, suggesting a synaptic vulnerability associated with AMPA receptor subunit imbalance in postnatal development that resulted in functional impairment of the mature cochlea.

A non-invasive functional assay of cochlear afferent transmission is wave-1 of the ABR, reflecting synchronous action potential generation in the auditory nerve at the onset of sound. With louder sounds, ABR wave-1 becomes larger. As synapses are lost or the spikes they evoke become less synchronous, ABR wave-1 becomes smaller. In adult mice, females have larger wave-1 amplitudes than males (Lozier et al., 2023; Milon et al., 2018; Rouse et al., 2020). Also, female synapses appear to be more resistant to noise-induced hearing loss, and their wave-1 amplitudes are more resistant to reduction in size following noise exposure (Milon et al., 2018; Shuster et al., 2021); these sex differences seem related to estradiol. However, the effector molecules mediating these differences have yet to be discovered. Analysis of sex-dependent mRNA expression in the mouse cochlea suggests that *Gria2-4* levels were similar overall but that females had more neurons with higher *Gria3* mRNA relative to *Gria2* and a greater abundance of fast-gating *flop* isoforms (Lozier et al., 2023). Interestingly, in humans and rodents, *Gria3* is located on the X-chromosome (Gécz et al., 1999; Li and Carrel, 2008), where, at least in fibroblasts, the gene is subject to X-inactivation. Here, we studied adolescent and mature female GluA3^KO^ mice raised in an ambient or low-level sound vivarium. We find that female mice are more sensitive to the absence of GluA3 than male mice. Indeed, raising female mice in a vivarium with normal ambient levels of sound resulted in early cochlear synaptopathy and hearing loss, whereas this effect was prevented by rearing in a low-level sound vivarium. In ambient sound, like males, cochlear synapses in female *Gria3^KO^* mice had less GluA2 and more GluA4 relative to *Gria3^WT^*. Thus, cochlear synapses rely on GluA3 subunits for proper structure and function, and we conclude that females depend on GluA3 to a greater degree than males.

## Results

### Cochlear output is reduced in 5-week-old GluA3^KO^ female mice raised in ambient sound but unaltered when raised in quiet

We performed a longitudinal study from P20 to P90 to determine the role of GluA3 in the hearing sensitivity of female mice. Statistical tests included two-way ANOVA for the comparisons of the click and pure tone thresholds and two-way mixed ANOVA followed by *post hoc* Holm-Sℒidák’s for wave 1 amplitude and latency. First, GluA3^WT^ and GluA3^KO^ mice were raised to P20 in very low-level sound (“quiet,” 10 dB SPL for mouse hearing; See Methods; **Fig. 1A**) compared to the normal vivarium ambient sound (40 dB SPL for mouse hearing). Data showed no difference between genotypes in ABR thresholds or wave-1 amplitudes at P20 (clicks p = 0.10, pure tones p > 0.05; wave-1 amplitude p > 0.05) (**Dataset EV1**). After verifying the two groups of mice not being different at baseline, we split the mice into two groups: ambient and quiet up to P90, to see if the sound level made a difference in the hearing sensitivity in the absence of GluA3 (**Fig. 1A**).

**Figure 1.**
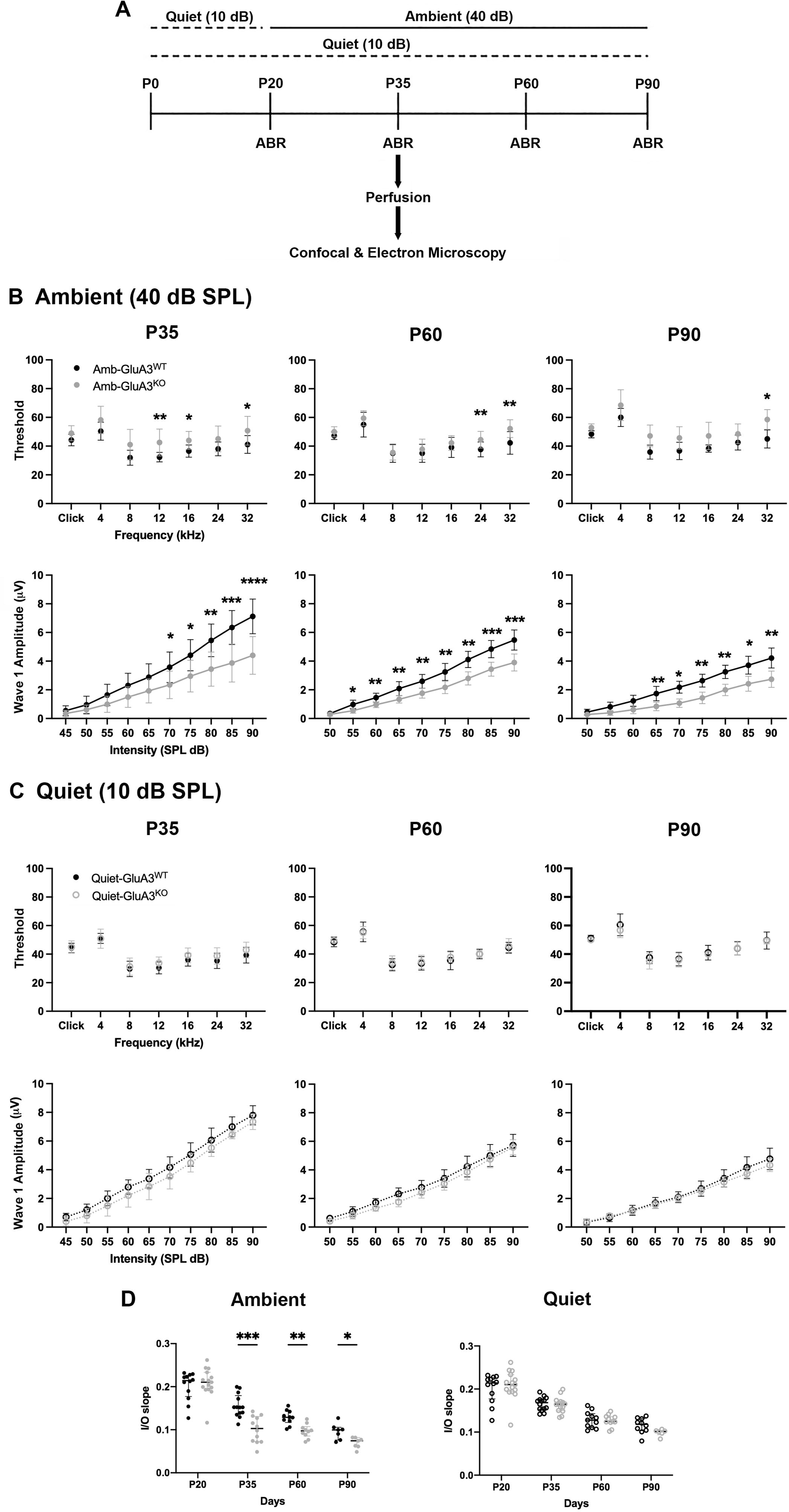
ABR wave-I amplitude was reduced, and threshold was elevated in female mice raised in ambient sound. **A)** Summary of the experimental longitudinal design: ABRs were recorded at four different time points from the mice raised either in a facility with quiet sound conditions (10 dB SPL; dotted line) or ambient sound (40 dB SPL; solid line). At P35, some of the GluA3^WT^ and GluA3^KO^ mice (n = 17) were transcardially perfused, and cochleae were collected for further ultrastructural and immunofluorescence, and confocal analyses. **B)** GluA3^WT^ and GluA3^KO^ mice raised in ambient sound levels. ABR data (mean ± SD); statistical tests included two-way ANOVA for the comparisons of the clicks and pure tone thresholds, and two-way mixed ANOVA followed by *post hoc* Holm-Sℒidák’s for wave-1 amplitude. **P35** (n =13 GluA3^WT^, n = 14 GluA3^KO^): click thresholds (p = 0.10), pure tones thresholds (p_12kHz_, p_16kHz_ and p_32kHz_ < 0.05; p_4kHz_, p_8kHz_ and p_24kHz_ p > 0.05), wave-1 amplitude above 65 dB SPL (p < 0.05). **P60** (n = 10 for each genotype): click thresholds (p = 0.75), pure tones thresholds (p_4k-16 kHz_ > 0.05; p_24-32kHz_ < 0.05) and wave-1 amplitude above 50 dB SPL (p < 0.05). **P90** (n=10 for each genotype): click thresholds (p = 0.07), pure tones thresholds (p_4k-24 kHz_ > 0.05, p_32 kHz_ < 0.05) and wave-1 amplitude above 60 dB SPL (p < 0.05). Asterisks indicate the statistical differences (*p < 0.05, **p < 0.01, ***p < 0.001, ****p < 0.0001). See **Dataset EV2**. **C)** GluA3^WT^ and GluA3^KO^ mice reared in quiet sound levels. ABR data (mean ± SD), statistical tests included two-way ANOVA for the comparisons of the clicks and pure tone thresholds, and two-way mixed ANOVA followed by *post hoc* Holm-Sℒidák’s for wave-1 amplitude. P35 (n =14 for each genotype), P60 (n =12 GluA3^WT^, n = 11 GluA3^KO^) and P90 (n =12 GluA3^WT^, n = 6 GluA3^KO)^: click thresholds (p > 0.05), pure tones thresholds (p_8-32 kHz_ > 0.05) and wave-1 amplitude (p > 0.05). See **Dataset EV1**. **D)** Input/output (I/O) slope of ABR wave-1 amplitude in GluA3^WT^ and GluA3^KO^ from P20 to P90 in ambient and quiet. Two-way ANOVA followed by *post hoc* Sℒidák’s multiple comparison test. In mice raised in ambient and relative to GluA3^WT^ mice the I/O slope was significant at P35 (p = 0.0004), at P60 (p = 0.0018) and at P90 (p = 0.045) (left). In mice raised in quiet the I/O slope was similar at all ages tested (P30, P60 and P90 p > 0.05). Each data point shows individual ABR wave-I amplitude I/O slope as function of sound intensity. Data represents the median. Asterisks indicate the statistical differences (*p < 0.05, **p < 0.01, ***p < 0.001,). (P20, n = 13 GluA3^WT^, n = 17 GluA3^KO^; the number of mice for P30, P60 and P90 is as in Fig. 1B and C).

The first female mice group were raised in the same ambient sound levels (**Fig. 1B**) as males (Rutherford et al., 2023). See figure legend for p values of clicks and pure tones thresholds and wave-1 amplitude. Our data show that postnatal day 35 (P35) female GluA3^KO^ mice had significantly elevated pure tone thresholds at 12, 16, and 32 kHz (**Fig. 1B top left; Dataset EV2**). Although there was an insignificant trend toward elevated response thresholds for the rest of the stimuli tested in response to clicks and 4, 8, and 24 kHz tones in the KO. In contrast to the male GluA3^KO^ mice (Rutherford et al., 2023), the female GluA3^KO^ mice had significantly smaller click wave-1 amplitudes above 65 dB SPL (**Fig. 1B bottom left; Dataset EV2**). To determine if reduced cochlear sensitivity in female GluA3^KO^ persisted, we further tested the mice at 2 months (P60) and 3 months (P90) of age, with similar findings. At P60, ABRs show that mice raised in ambient sound levels have similar click thresholds between genotypes, however, the GluA3^KO^ had elevated pure tone thresholds at 24 and 32 kHz, and reduced wave-1 amplitude above 50 dB SPL (**Fig. 1B top and bottom center; Dataset EV2**). At P90, ABR thresholds were significantly elevated in female KO mice at 32 kHz only, and wave-1 amplitude was significantly reduced above 60 dB SPL (**Fig. 1B top and bottom right; Dataset EV2**). The lack of significant threshold shifts at many frequencies at P90 despite the larger trends relative to P60 appeared to result from greater variance, potentially due to effects of the *Cdh23* age-related hearing loss phenotype in C57BL/6 mice. Wave-1 latency as a function of click level was similar between GluA3^KO^ and GluA3^WT^ at all ages tested (P20, P60, P90 p > 0.05; **Dataset EV2**). Our findings show that lack of the AMPA receptor subunit GluA3 accelerates hearing loss of females raised in ambient sound levels.

Next, we aged the GluA3^WT^ and GluA3^KO^ mice up to three months in the quiet and performed ABRs at P35, P60 and P90 (**Fig. 1C**). See figure legend for p values of clicks and pure tones thresholds and wave-1 amplitude. In contrast to mice raised in the normal vivarium, there was no difference between genotypes in ABR click threshold, pure-tone thresholds, or click wave-1 amplitude at any age for mice raised in quiet (**Fig. 1C top and bottom; Dataset EV1**). Wave-1 latency was also similar between the GluA3^WT^ and GluA3^KO^ mice at all ages tested (P20, P60, P90 p > 0.05; **Dataset EV1**).

We next analyzed wave-1 amplitude as a function of stimulus level (input/output (I/O) slope; two-way ANOVA followed by *post hoc* Sℒidák’s multiple comparison test; see figure legend for p values) between the two genotypes from P20 to P90 in ambient and in quiet (**Fig. 1D**). In ambient sound, we found that I/O slope at P35, P60, and P90 was significantly lower in GluA3^KO^ than GluA3^WT^ (**Fig. 1D left**). In contrast, we did not observe any difference in I/O slope between genotypes in quiet conditions (**Fig. 1D right**). As expected, I/O slope decreased with cochlear maturation and subsequent aging (Abdala and Dhar, 2012; Konrad-Martin et al., 2012; Johannesen and Lopez-Poveda, 2021). Interestingly, the largest reduction in I/O slope occurred for GluA3^KO^ mice raised in ambient sound between P20 to P35 (**Fig. 1D**), suggesting an ambient sound-evoked activity-dependent requirement for GluA3 during maturation of the cochlea. This effect on wave-1 amplitude suggests poor synchrony of action potential generation in the auditory nerve, potentially caused by synaptopathy in P35 GluA3^KO^ female mice raised in ambient conditions.

### Ambient sound triggers hearing loss in female GluA3^KO^ mice

As described above, raising the mice in ambient sound levels from P20 to P90 elevated cochlear thresholds and reduced ABR wave-1 in the auditory nerve. Figure 2 compares ABR thresholds and wave-1 amplitudes for WT or KO in ambient vs. quiet conditions at P35, P60 and P90. Statistical tests included two-way ANOVA for the comparisons of the click and pure tone thresholds and two-way mixed ANOVA for wave 1 amplitude followed by *post hoc* Holm-Sℒidák’s multiple comparison test at each age (see figure legend for p values). In GluA3^WT^ female mice we did not observe any differences in click and pure-tone thresholds, or wave-1 amplitudes when comparing mice raised in ambient sound to mice raised in low-level (i.e., quiet) sound at any age (**Fig. 2A**; **Dataset EV1 and EV2**). In GluA3^KO^ mice, click and pure-tone thresholds were similar at P35, except the 8 and 12 kHz pure tone thresholds which were significantly higher in the GluA3^KO^ mice raised in ambient sound compared to those in quiet (**Fig. 2B**; **Dataset EV1 and EV2**). At P60 clicks and pure tone thresholds were similar in GluA3^KO^ mice reared in ambient and in quiet except at 32 kHz thresholds that were higher in the KO mice raised in ambient. At P90, click thresholds were similar and the pure tone thresholds of the KO appeared to be higher in the ambient when compared to KO mice raised in quiet, but the difference was only statistically significant at 8 kHz (**Fig. 2B upper**). In contrast, wave-1 amplitude was significantly lower at all ages in GluA3^KO^ mice raised in ambient compared to those in quiet. The reduction in wave 1 amplitude in the KO in ambient was found significant above 65 dB SPL at P35, above 55 dB SPL at P60 and above 50 dB SPL at P90 (**Fig. 2B bottom**).

**Figure 2.**
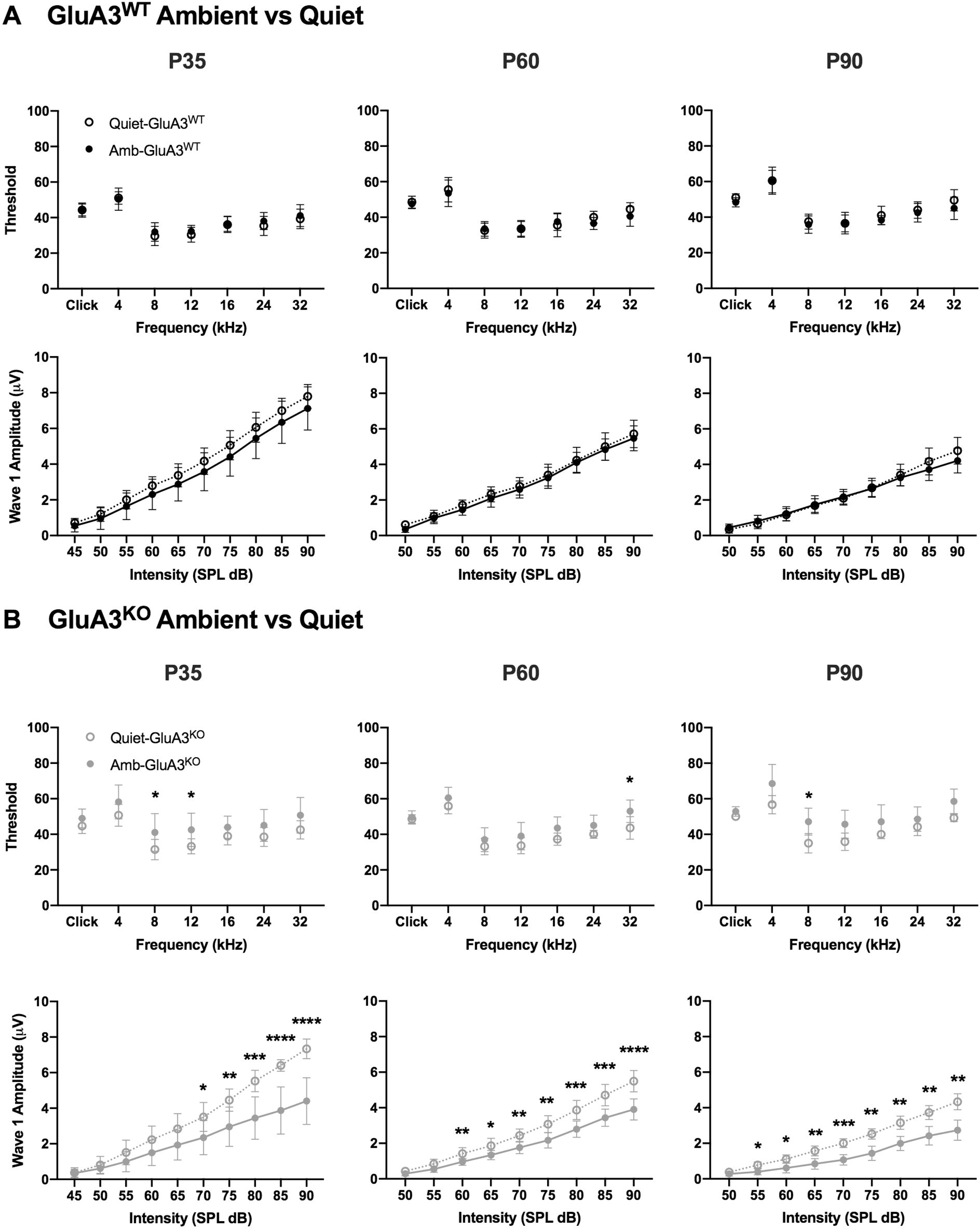
Ambient sound triggers ABR click wave-I amplitude reduction in female GluA3^KO^ mice. **A-B)** Statistical tests included two-way ANOVA for the comparisons of the click and pure tone thresholds and two-way mixed ANOVA for wave-1 amplitude followed by *post hoc* Holm-Sdák’s multiple at each age. **A)** GluA3^WT^ mice raised in ambient and quiet show similar ABR clicks and pure tones thresholds (upper panel), and click wave-1 amplitude (lower panel) at P35, P60 and P90 (clicks p > 0.05, pure tones p p_4-32 kHz_ > 0.05; wave-1 amplitude p > 0.05) for each of the age group). Data represented as mean ± SD. **B)** GluA3^KO^ mice raised in ambient and in quiet. ABR clicks and pure tones thresholds (upper panel), and click wave-I amplitude (lower panel) at P35, P60, and P90. **P35**: clicks thresholds (p = 0.16), pure tones thresholds (p_4 kHz_, p_16-32 kHz_ > 0.05, p_8 kHz_ < 0.048, p_12 kHz_ = 0.022) and wave-1 amplitude above 65 dB SPL (p < 0.05 to 0.0001). **P60**: click thresholds (p = 0.99), pure tones thresholds (p_4-24 kHz_ > 0.05, p_32 kHz_ = 0.022) and wave-1 amplitude above 55 dB SPL (p < 0.05 – 0.0001). **P90**: click thresholds (p = 0.19), pure tones threshold (p_4 kHz_, p_12-32 kHz_ > 0.05, p_8 kHz_ = 0.046) and wave-1 amplitude (p < 0.05 - 0.001). Asterisks: *p < 0.05, **p < 0.01, ***p < 0.001, ****p < 0.0001. Data represented as mean ± SD.

In mice, the first postnatal month is a sensitive and critical period for the development of inner hair cell (IHC) ribbon synapses and cochlear function (Rutherford and Moser, 2016; Wong et al., 2014; Michalski et al., 2019). After the onset of hearing function at the end of the 2^nd^ postnatal week, cochlear function continues to develop with the maturation of afferent axonal and synaptic anatomies (Wong et al., 2013; 2014; Kim and Rutherford, 2016; Payne et al., 2021). Next, we asked if this reduction in wave-1 amplitude in female mice lacking GluA3 and raised in ambient sound was associated with <u>cochlear synaptopathy</u>: the pathological alteration, degradation, or loss of ribbon synapses.

### Ambient sound triggers synaptopathy in female GluA3^KO^ mice

To assess cochlear synapses at 5 weeks of age (P35) with confocal microscopy, we immunolabelled the presynaptic ribbon and the postsynaptic AMPARs. Female GluA3^WT^ and GluA3^KO^ mice had similar numbers of paired synapses (juxtaposed pre- and post-synaptic markers) throughout the cochlea in both ambient and quiet conditions (**Fig. 3A left and right;** Two-tailed Mann-Whitney U test with Holm-Sℒidák’s correction for multiple comparisons for all tests in Figs. 3-4; see legends for exact *p* values ≥ 0.001). The number of paired synapses per IHC, mean across all images throughout the cochlea, was 15.3 and 15.5 for WT and KO, respectively in ambient sound (for means by cochlear region, see **Dataset EV3**). In quiet, the mean paired synapses per IHC were 17.3 in WT and 16.7 in KO. In contrast, the number of lone ribbons (anti-CtBP2 without juxtaposed postsynaptic markers) and the number of ribbonless synapses (anti-GluA2 and -GluA4 without juxtaposed CtBP2) per IHC were significantly greater throughout the cochlea in GluA3^KO^ compared to WT, but only in ambient sound (**Fig. 3B-C left**; lone ribbons: mean of 0.83 in WT vs. 2.4 in KO; ribbonless: 0.90 vs. 2.4). In quiet conditions, GluA3^KO^ had significantly fewer ribbonless synapses than WT in the 20 kHz region and when considering all regions as one (**Fig. 3C right;** mean of 1.1 in WT vs. 0.23 in KO), while lone ribbons in quiet were similar in WT and KO (**Fig. 3B right;** mean of 1.1 in WT vs. 0.8 in KO). Thus, relative to paired synapses (∼ 15 per IHC), lone ribbons and ribbonless synapses were relatively rare in quiet and in WT in ambient sound (∼ 0 – 2 per IHC, mean among images; group means of ∼ 1 per IHC). However, in GluA3^KO^ raised in ambient sound the group means were > 2 per IHC (**Fig. 3B-C left**).

**Figure 3.**
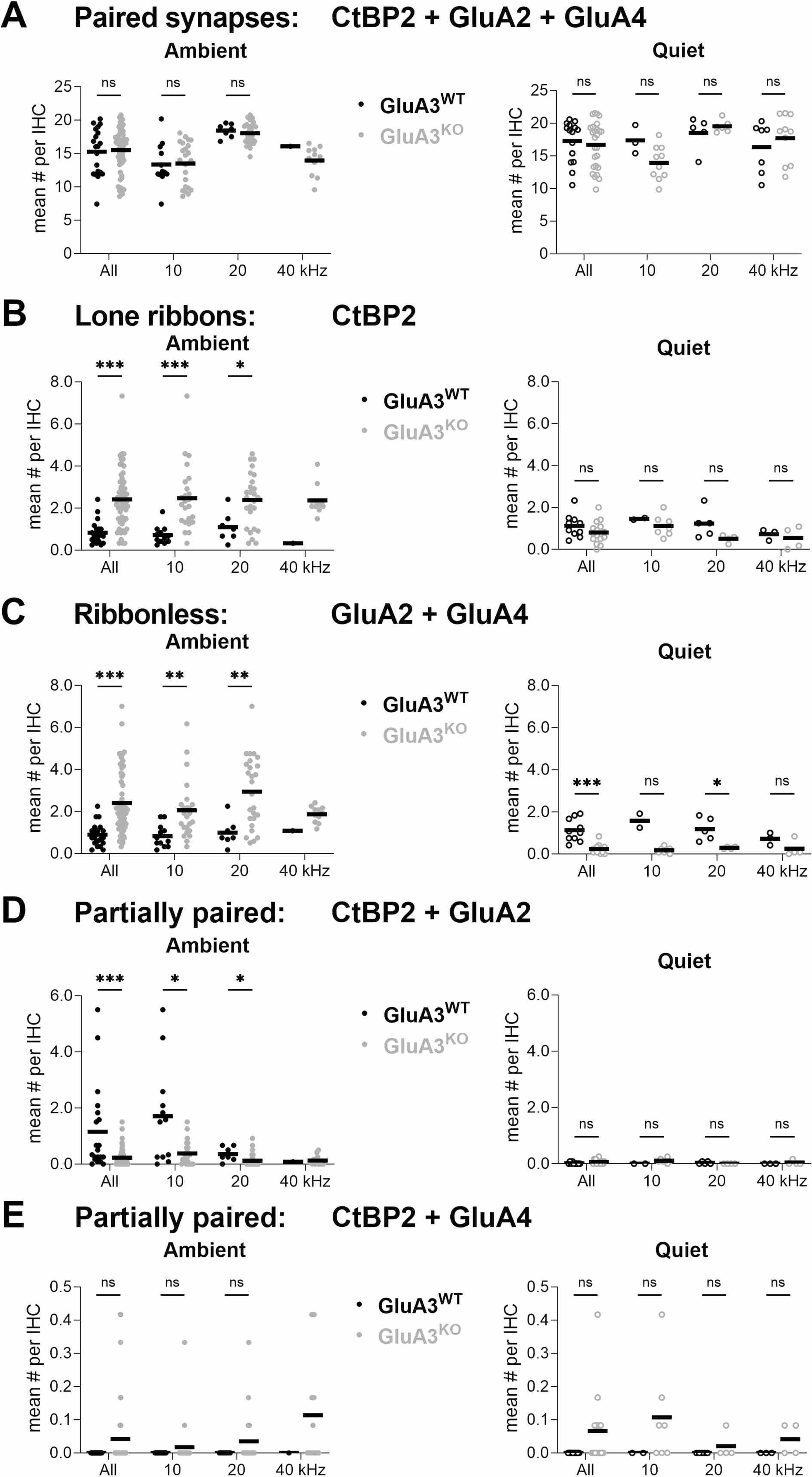
Synapse counts for 5-week-old females raised in ambient sound or in quiet. Number per inner hair cell for all synapses and divided into 3 tonotopic regions centered at 10, 20, or 40 kHz; GluA3^WT^ (black), GluA3^KO^ (gray); Ambient (left, filled circles, n = 20 WT and 61 KO images), Quiet (right, open circles, n = 15 WT and 26 KO images) from 4 WT and 5 KO mice. Markers are image means and bars are means of group means. Two-tailed Mann-Whitney U test with Holm-Sℒidák’s correction for multiple comparisons; ns p ≥ 0.05, * p < 0.05, ** p < 0.01, *** p < 0.001. The lowest reported p value is 0.001, so p values lower than 0.001 are reported as p < 0.001. **A)** Paired synapses (CtBP2 + GluA2 + GluA4) were not significantly different in number between GluA3^WT^ and GluA3^KO^ for all synapses or in any of the 3 tonotopic regions in ambient sound or in quiet. Ambient: Adjusted p = 0.90, 0.90, and 0.89 for all synapses, 10 kHz, and 20 kHz regions, respectively; for the 40 kHz region n = 1 image in WT precluded statistical comparison. Quiet: Adjusted P = 0.87, 0.27, 0.89, and 0.89 for all synapses, 10 kHz, 20 kHz, and 40 kHz regions, respectively **B)** Lone ribbons (CtBP2 only) were significantly greater in number in GluA3^KO^ than GluA3^WT^ in ambient sound; not significantly different in quiet. Ambient: Adjusted p = < 0.001, < 0.001, and 0.02 for all synapses, 10 kHz, and 20 kHz regions, respectively; for the 40 kHz region n = 1 image in WT precluded statistical comparison. Quiet: Adjusted p = 0.42, 0.44, 0.15, and 0.91 for all synapses, 10 kHz, 20 kHz, and 40 kHz regions, respectively **C)** Ribbonless synapses (GluA2 + GluA4) were significantly greater in number in GluA3^KO^ than GluA3^WT^ in ambient sound; but significantly fewer in quiet. Ambient: Adjusted p = < 0.001, 0.001, and 0.005 for all synapses, 10 kHz, and 20 kHz regions, respectively; for the 40 kHz region n = 1 image in WT precluded statistical comparison. Quiet: Adjusted p = < 0.001, 0.05, 0.04, and 0.17 for all synapses, 10 kHz, 20 kHz, and 40 kHz regions, respectively **D)** Synapses lacking detectable GluA4 (CtBP2 + GluA2 only) were significantly fewer in number in GluA3^KO^ than GluA3^WT^ in ambient sound; not significantly different in quiet. Ambient: Adjusted p = < 0.001, 0.03, and 0.03 for all synapses, 10 kHz, and 20 kHz regions, respectively; for the 40 kHz region n = 1 image in WT precluded statistical comparison. Quiet: Adjusted p = 0.42, 0.38, 0.69, and 0.99 for all synapses, 10 kHz, 20 kHz, and 40 kHz regions, respectively **E)** Synapses lacking detectable GluA2 (CtBP2 + GluA4 only) were sometimes observed, only in GluA3^KO^, but not significantly different in number in ambient sound or in quiet. Ambient: Adjusted p = 0.09, 0.54, and 0.51 for all synapses, 10 kHz, and 20 kHz regions, respectively; for the 40 kHz region n = 1 image in WT precluded statistical comparison. Quiet: Adjusted *p* = 0.08, 0.80, 0.80, and 0.80 for all synapses, 10 kHz, 20 kHz, and 40 kHz regions, respectively

**Figure 4.**
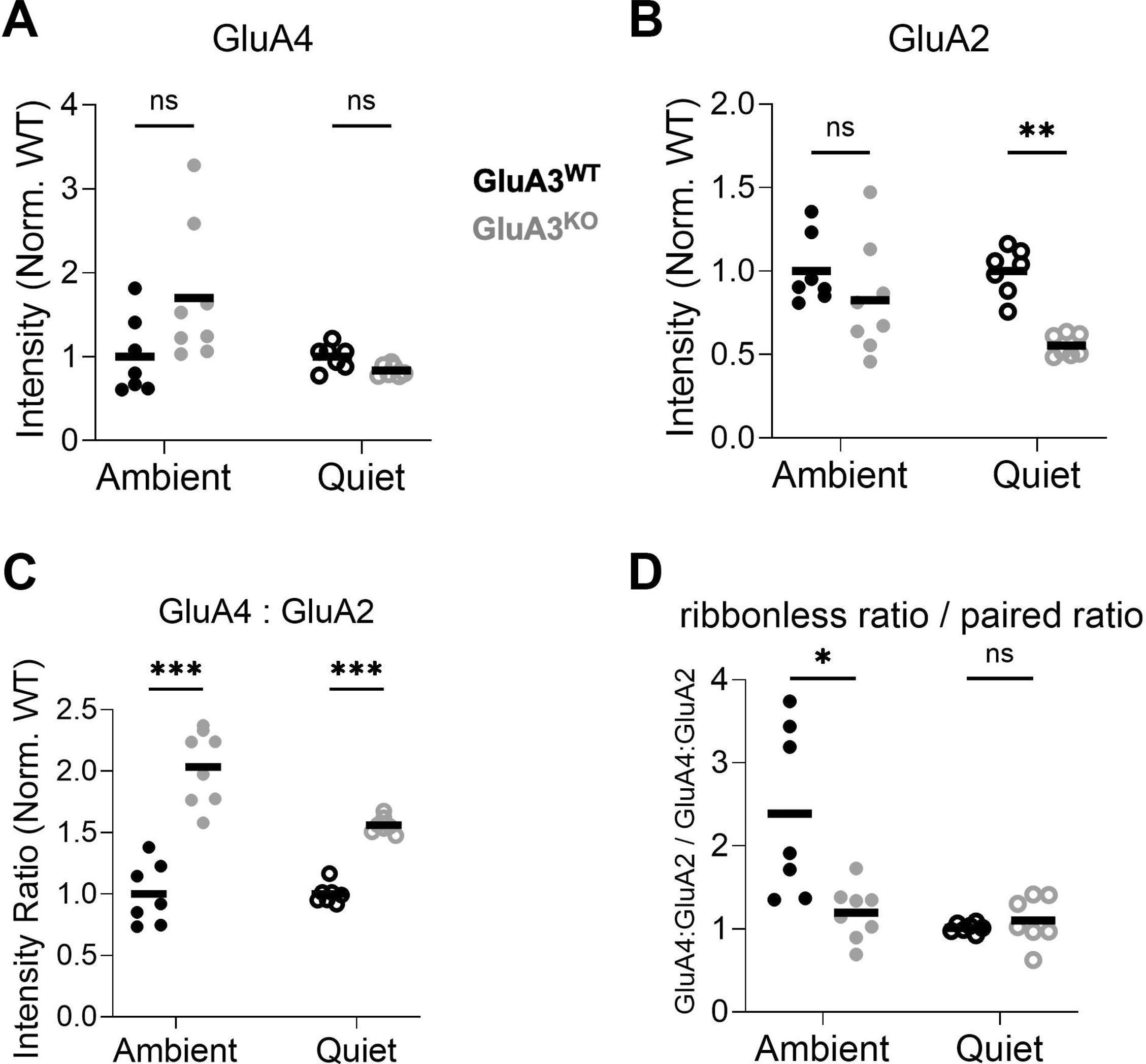
AMPAR intensity analysis of synapses from 5-week-old females raised in ambient sound or quiet. GluA3^WT^ (black), GluA3^KO^ (gray); Ambient (left, filled circles, n = 7 WT and 7 KO images), Quiet (right, open circles, n = 7 WT and 7 KO images). Markers are image means and bars are means of group means. Two-tailed Mann-Whitney U test with Holm-Sℒidák’s correction for multiple comparisons; ns p ≥ 0.05, * p < 0.05, ** p < 0.01, *** p < 0.001. Image means were normalized to the WT group mean per sound condition in panels A – C. **A)** For GluA4 intensity per synapse, there was no significant difference between genotypes in ambient (p = 0.07) or in quiet conditions (p = 0.07). **B)** For GluA2 intensity per synapse, there was no significant difference between genotypes in ambient (p = 0.15), but GluA3^WT^ synapses had significantly more GluA2 fluorescence in quiet conditions (p = 0.001). **C)** The GluA4:GluA2 intensity ratio per synapse was significantly greater for GluA3^WT^ synapses in both ambient and in quiet conditions (p = < 0.001). **D)** In each image, the mean GluA4:GluA2 intensity ratio for ribbonless synapses was divided by the mean GluA4:GluA2 intensity ratio for paired synapses, such that numbers greater than 1 indicate greater GluA4 fluorescence at ribbonless synapses than paired synapses. Ribbonless synapses had greater GluA4:GluA2 ratios than paired synapses for GluA3^WT^ mice in ambient conditions (p = 0.01) but they were similar in quiet (p = 0.8).

Some synapses (partially-paired) contained a presynaptic ribbon juxtaposed to only GluA2 or, very rarely, only GluA4. In ambient sound, GluA3^KO^ had significantly fewer GluA2-only synapses per IHC than WT throughout the cochlea (**Fig. 3D left;** mean of 1.2 in WT vs. 0.23 in KO; for means by cochlear region, see **Dataset EV3**) and an insignificant trend toward a greater number of GluA4-only synapses than WT (**Fig. 3E left;** mean of 0.0 in WT vs. 0.042 in per IHC in KO). In quiet, both varieties of partially-paired synapses were very rare: GluA2-only synapses were rarely seen in either genotype (**Fig. 3D right;** mean of 0.02 in WT vs. 0.06 per IHC in KO), while GluA4-only synapses were seen only in GluA3^KO^ (**Fig. 3E right;** mean of 0 in WT vs. 0.07 per IHC in KO). Thus, GluA4-only synapses were seen only in GluA3^KO^ for females in ambient or quiet conditions. Although GluA4-only synapses were significantly greater in number in GluA3^KO^ males than WT raised in ambient conditions (Rutherford et al., 2023), the differences did not reach statistical significance for the females (All synapses ambient, **Fig. 3E left**; All synapses quiet, **Fig. 3E right**).

Next, we compared synapse size and protein expression based on immunofluorescence volume and intensity per synapse, displayed as means per image and per group after normalizing metrics from individual images to the respective WT mean for images from ambient or quiet conditions (**Fig. 4**). Experiments were performed with GluA3^KO^ and GluA3^WT^ in parallel, separately for the two sound conditions, so comparisons are made only between GluA3^WT^ and GluA3^KO^. The overall volume of paired synapses appeared significantly smaller in GluA3^KO^ in quiet compared to GluA3^WT^ (11% smaller than WT; p = 0.03; not shown); the two genotypes were similar in ambient sound (2.2% smaller than WT; p = 0.78; not shown). CtBP2 intensity did not differ by genotype in either condition (p = 0.54 in ambient, p = 0.10 in quiet). For GluA4 and GluA2 intensity per synapse (**Fig. 4A-B; Dataset EV4**), in quiet conditions, the variance between images was relatively small, and the decrease in GluA2 intensity in the GluA3^KO^ was significant (45% less than WT; **Fig. 4B**; see legend for exact p values). For GluA4 and GluA2 fluorescence intensity per synapse in ambient sound, we observed the same trends as in the males: an increase in GluA4 and a decrease in GluA2 in GluA3^KO^. Although these two trends did not reach statistical significance in ambient sound due to increased variance (**Fig. 4A-B**), the GluA4:GluA2 ratios were significantly increased in both quiet and ambient sound (103% increase in ambient vs. 56% increase in quiet (**Fig. 4C**). We next asked if synapses without a ribbon differed in GluA4:GluA2 ratio from synapses with a ribbon by comparing the ratios between those two groups of synapses within each image. Only in GluA3^WT^ mice in ambient conditions there was an obvious difference between ribbonnless synapses and paired synapses, where the GluA4:GluA2 ratio was greater at ribbonless synapses than paired synapses (**Fig. 4D**). As well, in ambient sound the ribbonless ratio / paired ratio was significantly greater in WT than KO by 99%.

At 5 weeks, the female GluA3^KO^ mice raised in quiet had apparently normal ABR (**Fig. 1-2**) and numbers of paired synapses (**Fig. 3**). Still, they had significantly reduced expression of GluA2 per synapse (**Fig. 4**). Next, we looked within synapses for additional anatomical synaptopathy. Similar to a previous study (Hu et al., 2020), we selected well-isolated synapses and divided each synapse into nanodomains to measure the GluA4:GluA2 fluorescence ratio (**Fig. 5A-B**; **Fig. EV1;** see Methods). In both genotypes, we observed a broad range of ratios within nanodomains, generally between ∼ 0.1 – 2.0. However, in GluA3^KO^, the ratios for whole synapses and nanodomains extended to greater values than in GluA3^WT^, suggesting a greater fraction of AMPARs lacking the subunit GluA2 in GluA3^KO^ (**Fig. 5C-E**). For whole synapses in WT: mean ratio 0.58 ± 0.13 (S.D.), range 0.29 – 0.95; in KO: mean ratio 0.76 ± 0.23, range 0.31 – 1.42. For nanodomains in WT: mean ratio 0.53 ± 0.24, range 0.099 – 1.77; in KO: mean ratio 0.70, range 0.12 – 2.8. Indeed, the probability distributions for synapses and nanodomains differed between GluA3^WT^ and GluA3^KO^ (KS test: synapses, p = 7.2 e^-9^; nanodomains, p = 3.5 e^-19^). The positive relationship between GluA4:GluA2 ratio and ribbon fluorescence per synapse is shown in **Figure 5F**, and the strength of the correlation was tested with Spearman’s coefficient r_s_. For GluA3^WT^, the correlation was significant (r_s_ 0.26 > critical value 0.18); for GluA3^KO^, the correlation was not significant (r_s_ 0.092 < critical value 0.18).

**Figure 5.**
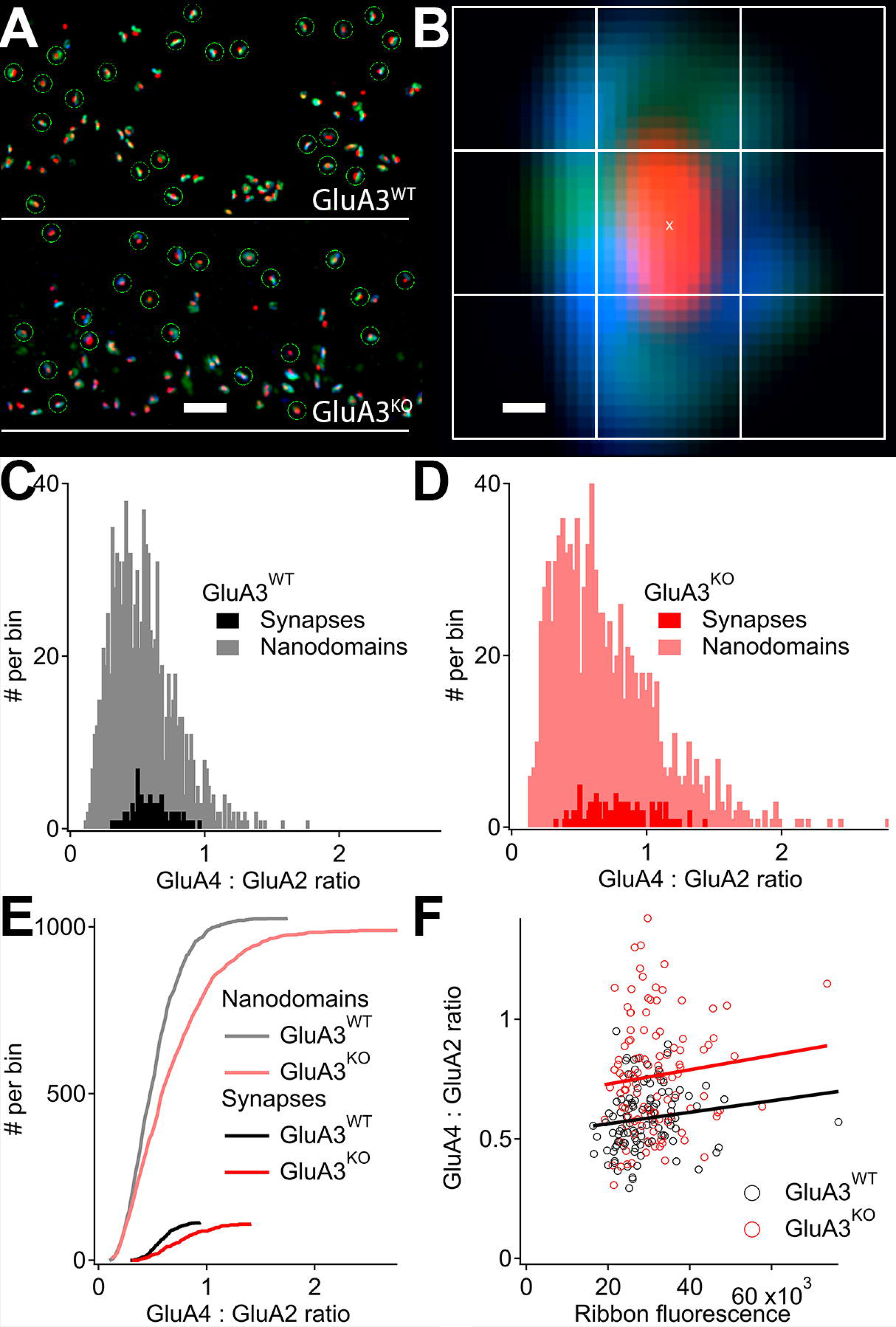
AMPA receptor subunit fluorescence ratios in whole synapses and nanodomains from 5-week-old female mice raised in quiet. **A)** GluA3^WT^ (upper) and GluA3^KO^ (lower) synapses on inner hair cells in the 38 kHz tonotopic region labeled with anti-CtBP2 (red), anti-GluA4 (blue), and anti-GluA2 (green). **B)** Individual synapse with 9 square 13×13 pixel matrices overlaid for nanodomain analysis. Each pixel is 26 nm. **C-D)** Frequency histograms of ratios of raw GluA4:GluA2 fluorescence for whole synapses (black, dark red) and nanodomains (gray, light red) for GluA3^WT^ (C) and GluA3^KO^ (D). Data from midcochlea. WT: 1,026 nanodomains in 114 synapses from 4 images; KO: 990 nanodomains in 110 synapses from 4 images. **E)** Cumulative histograms of the data in C-D. Comparing genotypes, the distributions of ratios were significantly different for synapses (KS test, p = 7.2 e^-9^) and for nanodomains within synapses (nanodomains, p = 3.5 e^-19^). **F)** GluA4:GluA2 fluorescence ratios per synapse as a function of ribbon fluorescence. The variables were significantly correlated for GluA3^WT^ (r_s_ 0.26 > critical value 0.18) but not for GluA3^KO^ (r_s_ 0.092 < critical value 0.18). Scale bars: A) 2 µm; B) 100 nm.

In the same set of synapses from female mice raised in quiet, we asked if the GluA4:GluA2 fluorescence ratios differed as a function of distance from the center of the synapse with radial distributions (**Fig. 6**; **Fig. EV1;** see Methods). Shells of pixel areas were created from concentric squares overlaid on maximum-intensity projections of Airyscan images in GluA3^WT^ and KO (**Fig. 6A-B**). For each fluorescence channel in each shell, the sum of pixel intensities was divided by the number of pixels to calculate shell density, and the GluA4:GluA2 density ratios were calculated for each shell in each synapse based on the raw values or the peak-normalized values (**Fig. 6C-D**). Radial distributions of GluA4:GluA2 fluorescence ratio per synapse were grouped by genotype in 6 tonotopic regions (**Fig. 6E-F**). This analysis confirmed the observation of greater GluA4:GluA2 fluorescence ratios in GluA3^KO^ relative to WT and shows this to be the case throughout intrasynaptic space when comparing similar tonotopic regions. On average, ratios tended to peak 200 – 400 nm from the synapse center in both genotypes, but this peak was less pronounced in GluA3^KO^ because GluA4:GluA2 ratios tended to be greater near the synapse center in GluA3^KO^ relative to GluA3^WT^. Thus, it appears that lack of GluA3 alters the intrasynaptic distribution of GluA2 and GluA4.

**Figure 6.**
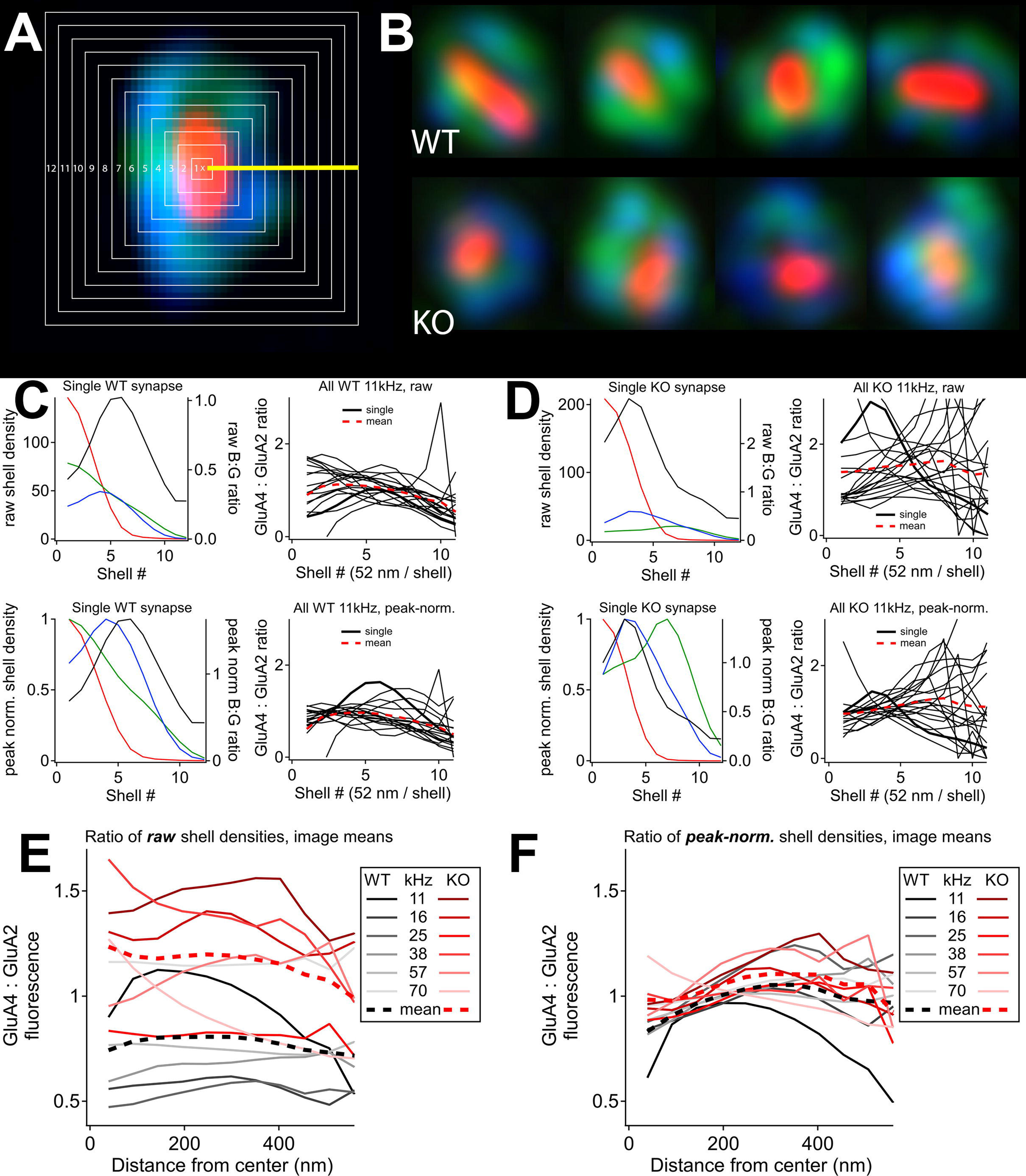
Radial distribution of AMPA receptors around synapse center in 5-week-old female mice raised in quiet. **A)** Squares were overlaid on synapses in 2-dimensional maximum-intensity projections to create shells for calculation of 1-dimensional radial distributions of blue to green fluorescence density to approximate changes in the relative abundance of GluA4:GluA2 as a function of distance from ribbon center. **B)** Examples of synapses from GluA3^WT^ (upper) and GluA3^KO^ (lower). **C)** Upper left: Radial distribution of raw shell densities for the WT synapse in (A) for CtBP2 (red), GluA4 (blue), and GluA2 (green). In black is the ratio of raw blue to green fluorescence. Upper right: Ratio of raw GluA4:GluA2 shell densities for the WT synapses from the image of the 11 kHz region; single synapse on the left is shown in thick black trace; the image mean is shown in the dashed red trace. Lower left: Radial distribution for the same synapse when normalized to the peak shell density of each fluorescence channel. Lower right: Ratio of peak-normalized GluA4:GluA2 shell densities for the same image. **D)** Same as (C) for the KO image from the 11 kHz region. **E)** Image means of the ratio of raw shell densities for WT and KO. Group means are in thick dashed lines for WT (black) and KO (red). **F)** Same as (E) for the ratio of peak-normalized shell densities

### Afferent swellings in the 5-week-old female GluA3^KO^ in ambient

To look for structural correlates of hearing impairment in the GluA3-deficient cochlea, we analyzed ultrastructurally the sensory epithelium of the mice raised in ambient and quiet sound conditions. The general structure and cellular components of the sensory epithelium of the female GluA3^KO^ raised in ambient and quiet were similar to GluA3^WT^ and published data of C57BL/6 mice (e.g., Ohlemiller and Gagnon, 2004). However, in the P35 female KO raised in ambient conditions, we noticed the existence of vacuoles underneath IHCs, which were most often observed in the high to middle frequency, basal-middle part of the cochlear spiral (**Fig. 7D, F, G; Fig. EV2**). We confirmed that the vacuoles corresponded to swellings of Type I afferent terminals synapsing on both sides of the IHCs (modiolar and pillar, **Fig. 7F, G; Fig. EV2**). Relative to other parts of the neuronal cytoplasm, these vacuoles were clearer, suggesting less density of electron-absorbing intracellular material. Some afferent swellings had a post-synaptic density (PSD) juxtaposed with a membrane-anchored pre-synaptic ribbon (**Fig. 7F)**, whereas others were ribbonless (**Fig. 7G**). The cytoplasm of the swollen afferents showed missing mitochondria and swollen endoplasmic reticulum. Alongside the swollen afferents, normal-looking afferent terminals with electron-dense PSDs and membrane-anchored presynaptic ribbons were present (**Fig. 7E**). We calculated the average gray value (AGV) of the cytoplasm of the afferent terminals and found that in the GluA3^KO^ the cytoplasm of the swollen afferents (AGV mean = 150 ± 23.4 ±SD; AGV scale settings black= 0 and white= 255) was significantly lighter than the “normal-looking” afferents in the KO (AGV mean = 105 ± 20.2 ±SD) or the GluA3^WT^ afferents (AGV mean = 102 ± 11.6 ±SD) (p< 0.0001, one-way ANOVA). A paired comparison between the AGV of the WT afferents and the “normal-looking” afferents in the GluA3^KO^ showed no difference (p > 0.005). In addition, we found that the swollen afferents in GluA3^KO^ (circularity mean = 0.8 ± 0.1 ± SD; roundness mean = 0.74 ± 0.13 ±SD) were significantly more circular and rounder than the “normal-looking” afferents in the KO (circularity mean = 0.6 ± 0.11 ± SD; roundness mean = 0.44 ± 0.15 ±SD) or the GluA3^WT^ afferents (circularity mean = 0.43 ± 0.1 ± SD; roundness mean = 0.41 ± 0.17 ±SD) (p < 0.0001, one-way ANOVA). A paired comparison between the “normal-looking” afferents in the KO and the GluA3^WT^ afferents showed that the formers were more circular (p = 0.045). In this analysis, we did not separate the data between modiolar and pillar side synapses. The observation of vacuole-associated Type I afferent swellings increased in the female KO at P60 (**Fig. EV2)** and were also observed in the apex of the cochlear spiral. Swelling of afferent terminals suggests glutamate excitotoxicity and synaptophathy in GluA3^KO^, reminiscent of noise-exposed or ischemic tissue from animals with intact GluA3 (Robertson 1983; Puel et al., 1994, 1998). Vacuoles or afferent swellings were not observed in female GluA3^WT^ mice in ambient or in either genotype raised in quiet, in which only normal-looking synapses were observed (**Fig. 7A-C, H-I, J-K**). Our results show chronic synaptopathy appearing ultra-structurally and physiologically before potential deafferentation of the IHC in female GluA3^KO^ mice raised in ambient sound.

**Figure 7.**
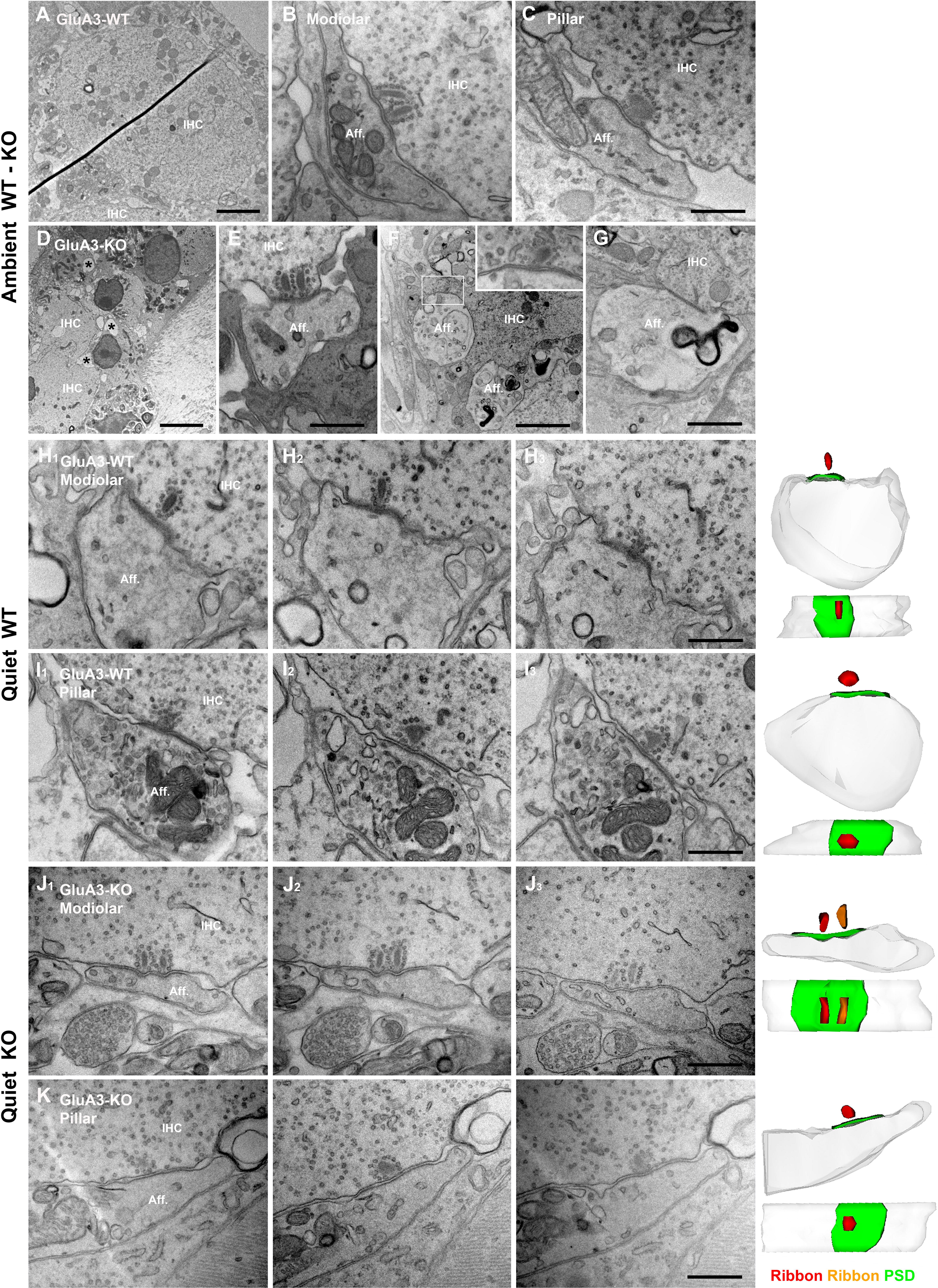
Pre- and post-synaptic ultrastructural features of IHC-ribbon synapses of 5-week-old female GluA3^WT^ and GluA3^KO^ mice. **A-C)** Representative transmission electron microscopy (TEM) micrographs of IHC-ribbon synapses of GluA3^WT^ mice (n = 2) raised in ambient sound. Micrographs are from the middle cochlea turn. **A)** The micrograph shows a low magnification of one inner hair cell (IHC) in cross-section. Scale bar: 2 μm. **B-C)** Micrographs show one modiolar (B) and one pillar (C) side IHC-ribbon synapse. The modiolar-side synapse has two presynaptic ribbons. Scale bar: 0.5 μm. Aff.: afferent. **D-G)** Representative TEM micrographs of IHC-ribbon synapses of GluA3^KO^ mice (n = 2) raised ambient sound. Micrographs are from the middle cochlea turn. **D)** TEM micrograph showing afferent swellings (asterisks) around IHCs. Scale bar: 5 μm. **E)** The micrograph shows a normal-looking double-ribbon modiolar IHC-ribbon synapse, Scale bar: 0.5 μm. **F)** Micrograph shows afferent terminal swellings synapsing on an IHC. Scale bar: 2 μm. The inset shows a higher magnification of the white box region. The insets show details of pre- and post-synaptic membranes with the adjacent presynaptic ribbon. **G)** Micrograph shows one afferent swelling with a flat postsynaptic membrane but without a pre-synaptic ribbon. Scale bar: 0.5 μm. **H_1_-I_3_)** Representative TEM serial electron micrographs of one IHC-ribbon synapse on the modiolar side (H_1_-H_3_) and on the pillar side (I_1_-I_3_) with their corresponding 3D reconstructions (right side) and its 3D reconstruction (right side) of GluA3^WT^ mice raised in quiet. Micrographs are from the middle cochlea turn. Scale bar: 0.5 μm. **J_1_-K_3_)** Representative TEM serial electron micrographs of one IHC-ribbon synapse on the modiolar side (J_1_-J_3_) and the pillar side (K_1_-K_3_) with their corresponding 3D reconstructions (right side) of GluA3^KO^ mice raised in quiet. No afferent swellings are observed in the GluA3^KO^ in the quiet. Micrographs are from the middle cochlea turn. Scale bar: 0.5 μm.

### Pre- and post-synaptic ultrastructural features of IHC-ribbon synapses in 5-week-old female mice raised in quiet

Previously, we reported that P35 male GluA3^KO^ mice raised in ambient sound levels had normal ABR and a similar number of paired IHC-ribbon synapses. However, those male KO mice showed defective differentiation of modiolar-side and pillar-side pre- and post-synaptic ultrastructural specializations (Rutherford et al., 2023). Due to the abundance of swollen afferents in the female GluA3^KO^ in ambient, we performed morphometric analysis of the pre- and post-synaptic structure of the modiolar- and pillar-side IHC-ribbon synapses in 5-week GluA3^WT^ and GluA3^KO^ mice raised in quiet. This analysis of synapses from mice raised in quiet allows us to determine structural alterations associated with increased synaptic vulnerability in louder sound environments.

### Ultrastructure in C57BL/6 GluA3^WT^ in quiet

Thirty-five IHC-ribbon synapses of female GluA3^WT^ mice were analyzed in three dimensions (3D) using ultrathin serial sections (on average, 10 ultrathin sections per PSD). Of this total, 24 synapses were on the modiolar side and 11 on the pillar side of IHCs. In our sample of modiolar-side synapses, 19 had one single ribbon, whereas 5 had two ribbons. All pillar-side synapses had only one ribbon. We did not find differences in PSD surface area or volume between the modiolar synapses with one or two ribbons, so data was combined. Next, we compared the PSD surface area and volume between modiolar side and pillar side and did not observe any difference in GluA3^WT^ (surface area p = 0.52; volume p = 0.10, unpaired t-test) (**Fig. 8A top; Table 1**). In contrast, male GluA3^WT^ PSDs were larger on the modiolar side (Rutherford et al., 2023). Data from single ultrathin sections showed that the PSD linear length was similar between the modiolar- and pillar-side synapses (p = 0.93, Mann-Whitney test) (**Fig. 8B top; Table 2**).

**Figure 8.**
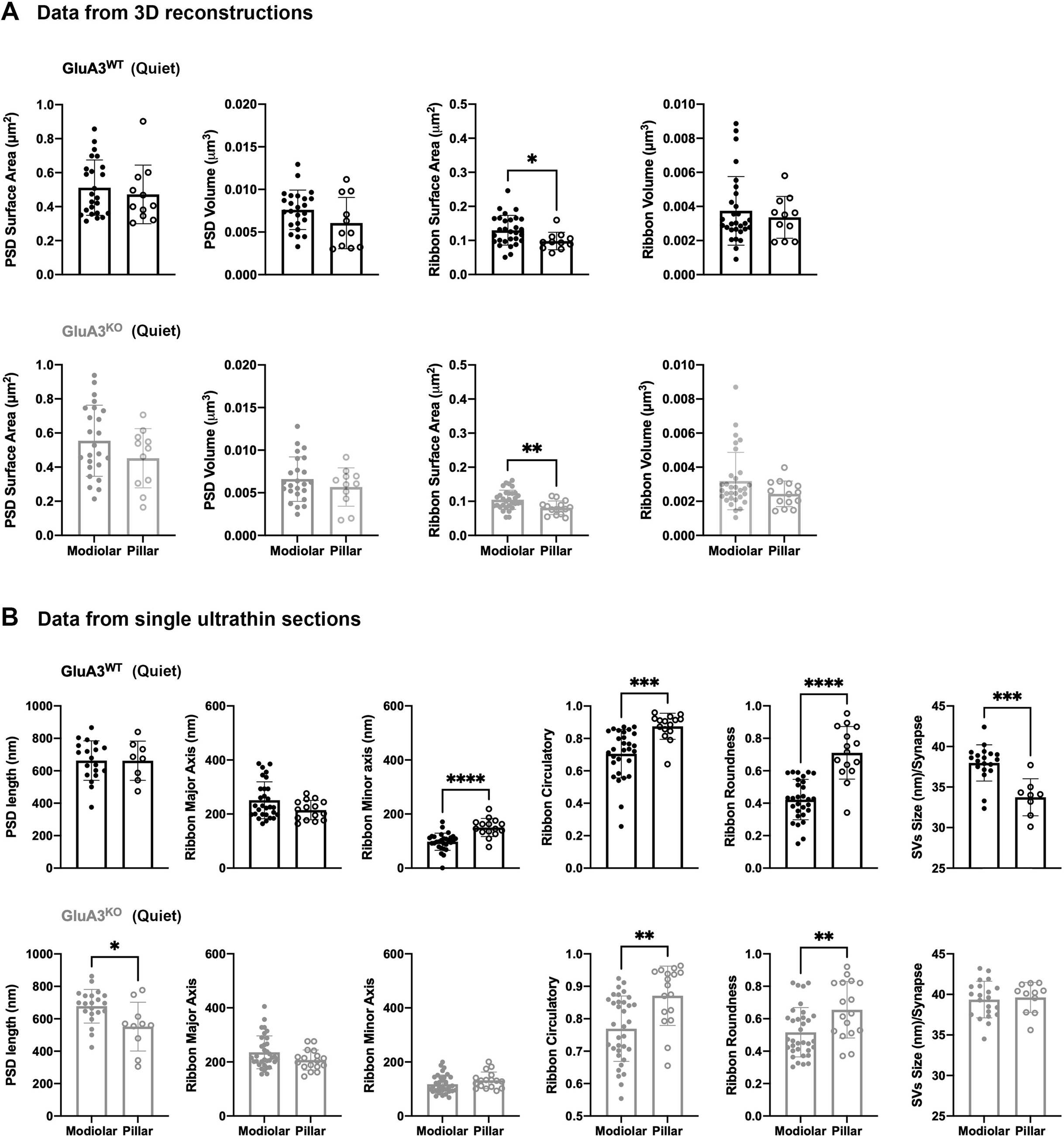
Morphometric analysis of mid-cochlear IHC-ribbon synapses of P35 GluA3^WT^ and GluA3^KO^ mice in quiet. **A)** Plots of the quantitative data from 3D reconstructions of the PSD and ribbon surface area and volume of GluA3^WT^ (black, upper panel) and GluA3^KO^ mice (gray, lower panel). Data represented as mean ± SD, unpaired t-test, *p < 0.05, **p < 0.01; n = 35 synapses per genotype. See Table 1 for descriptive analysis. **B)** Plots of the quantitative data from single ultrathin sections of the PSD length, ribbon major and minor axis, ribbon circularity and roundness, and synaptic vesicles size (SVs) of IHC-ribbon synapses of GluA3^WT^ (black, upper panel) and GluA3^KO^ (gray, lower panel). Data represented as mean ± SD, unpaired t-test, *p < 0.05, **p < 0.01, Mann-Whitney test; Synapses n = 62 GluA3^WT^, n = 66 GluA3^KO^. See Table 2 for descriptive analysis.

**Table 1.** Data from 3D reconstructions of IHC-ribbon synapses from mice raised in quiet.

| Genotype | Parameters | PSD surface area ($\mu\text{m}^2$) | | PSD volume ($\mu\text{m}^3$) | | Ribbon surface area ($\mu\text{m}^2$) | | Ribbon volume ($\mu\text{m}^3$) | |
| --- | --- | --- | --- | --- | --- | --- | --- | --- | --- |
|  |  | M | P | M | P | M | P | M | P |
| **GluA3<sup>WT</sup>** | Minimum | 0.32 | 0.30 | 0.003 | 0.003 | 0.05 | 0.06 | 0.001 | 0.002 |
|  | 25% Percentile | 0.36 | 0.35 | 0.006 | 0.003 | 0.10 | 0.08 | 0.003 | 0.002 |
|  | Median | 0.46 | 0.41 | 0.008 | 0.005 | 0.13 | 0.09 | 0.003 | 0.004 |
|  | 75% Percentile | 0.63 | 0.57 | 0.009 | 0.010 | 0.16 | 0.11 | 0.004 | 0.004 |
|  | Maximum | 0.86 | 0.90 | 0.013 | 0.011 | 0.25 | 0.16 | 0.009 | 0.006 |
|  | Mean | 0.51 | 0.47 | 0.008 | 0.006 | 0.13 | 0.10 | 0.004 | 0.003 |
|  | Std. Deviation | 0.16 | 0.17 | 0.002 | 0.003 | 0.04 | 0.03 | 0.002 | 0.001 |
|  | Coefficient of variation | 32% | 37% | 30% | 50% | 34% | 26% | 54% | 36% |
| **GluA3<sup>KO</sup>** | Minimum | 0.21 | 0.16 | 0.002 | 0.002 | 0.05 | 0.05 | 0.001 | 0.001 |
|  | 25% Percentile | 0.40 | 0.31 | 0.005 | 0.004 | 0.09 | 0.06 | 0.002 | 0.002 |
|  | Median | 0.50 | 0.51 | 0.006 | 0.006 | 0.10 | 0.08 | 0.003 | 0.002 |
|  | 75% Percentile | 0.74 | 0.58 | 0.009 | 0.007 | 0.12 | 0.09 | 0.004 | 0.003 |
|  | Maximum | 0.94 | 0.71 | 0.013 | 0.009 | 0.16 | 0.12 | 0.009 | 0.004 |
|  | Mean | 0.55 | 0.45 | 0.007 | 0.006 | 0.10 | 0.08 | 0.003 | 0.002 |
|  | Std. Deviation | 0.21 | 0.17 | 0.003 | 0.002 | 0.03 | 0.02 | 0.002 | 0.001 |
|  | Coefficient of variation | 38% | 38% | 40% | 40% | 27% | 25% | 53% | 31% |

M: Modiolar; P: Pillar

**Table 2.** Data from single ultrathin sections of IHC-ribbon synapses from mice raised in quiet.

| Genotype | Parameters | PSD length (nm) |  | Major axis (nm) |  | Minor axis (nm) |  | Circularity |  | Roundness |  | SVs size (nm) |  |
| --- | --- | --- | --- | --- | --- | --- | --- | --- | --- | --- | --- | --- | --- |
|  |  | M | P | M | P | M | P | M | P | M | P | M | P |
| **GluA3<sup>WT</sup>** | Minimum | 375 | 474 | 164 | 164 | 1 | 78 | 0.26 | 0.64 | 0.15 | 0.34 | 32 | 30 |
|  | 25% Percentile | 573 | 553 | 198 | 177 | 83 | 134 | 0.62 | 0.85 | 0.34 | 0.60 | 37 | 32 |
|  | Median | 679 | 680 | 226 | 217 | 99 | 146 | 0.73 | 0.91 | 0.43 | 0.72 | 38 | 34 |
|  | 75% Percentile | 756 | 754 | 293 | 244 | 114 | 169 | 0.83 | 0.92 | 0.54 | 0.87 | 39 | 35 |
|  | Maximum | 866 | 839 | 387 | 276 | 171 | 219 | 0.87 | 0.96 | 0.59 | 0.95 | 42 | 38 |
|  | Mean | 663 | 662 | 251 | 215 | 98 | 150 | 0.71 | 0.87 | 0.42 | 0.71 | 38 | 34 |
|  | Std. Deviation | 122 | 121 | 68 | 36 | 32 | 33 | 0.15 | 0.08 | 0.12 | 0.16 | 2 | 2 |
|  | Coefficient of variation | 18% | 18% | 27% | 17% | 33% | 22% | 21% | 9% | 30% | 23% | 6% | 7% |
| **GluA3<sup>KO</sup>** | Minimum | 424 | 305 | 155 | 146 | 68 | 93 | 0.55 | 0.66 | 0.30 | 0.37 | 35 | 36 |
|  | 25% Percentile | 635 | 446 | 198 | 182 | 92 | 105 | 0.70 | 0.81 | 0.40 | 0.52 | 38 | 38 |
|  | Median | 696 | 564 | 216 | 204 | 107 | 131 | 0.78 | 0.90 | 0.46 | 0.63 | 40 | 40 |
|  | 75% Percentile | 743 | 648 | 272 | 232 | 143 | 147 | 0.85 | 0.94 | 0.61 | 0.82 | 41 | 41 |
|  | Maximum | 863 | 777 | 405 | 276 | 200 | 201 | 0.92 | 0.96 | 0.82 | 0.92 | 43 | 41 |
|  | Mean | 678 | 552 | 236 | 207 | 118 | 132 | 0.77 | 0.87 | 0.52 | 0.66 | 39 | 40 |
|  | Std. Deviation | 104 | 150 | 61 | 37 | 34 | 31 | 0.10 | 0.09 | 0.15 | 0.17 | 2 | 2 |
|  | Coefficient of variation | 15% | 27% | 26% | 18% | 29% | 23% | 13% | 11% | 29% | 27% | 6% | 5% |

M: Modiolar; P: Pillar

We next compared the presynaptic ribbon surface area and volume between modiolar- and pillar-side synapses in GluA3^WT^ (**Fig. 8A top; Table 1**). The ribbon surface area of the modiolar-side was significantly larger than the pillar-side (p = 0.03, unpaired t-test). In contrast, ribbon volume was similar (p = 0.56).

Analysis of the presynaptic ribbon major and minor axes (unpaired t-test) showed that the major axis of modiolar-side ribbons was not significantly different from the pillar side (p = 0.06) (**Fig. 8B top; Table 2)**, in contrast to GluA3^WT^ males (Rutherford et al., 2023). The minor axis of modiolar-side ribbons was significantly smaller than the pillar side (p = <0.0001) (**Fig. 8B top; Table 2**). Moreover, ribbons of the modiolar-side were less circular (p = 0.0002, unpaired t-test) and more elliptical (p = 0.0001 unpaired t-test) than those on the pillar side (**Fig. 8B top; Table 2**), as was observed in the males. Analysis of synaptic vesicles (SVs) showed that those of the modiolar-side were larger than those of the pillar-side (p = 0.0005, Mann-Whitney test) (**Fig. 8B top; Table 2**), as in the males (Rutherford et al., 2023).

### Ultrastructure in C57BL/6 GluA3^KO^ in quiet

From GluA3^KO^, 35 IHC-ribbon synapses were analyzed in 3D with serial ultrathin sections (on average, 10 ultrathin sections per PSD). Of this total, 24 were on the modiolar side and 11 on the pillar side of the IHCs. Seventeen synapses on the modiolar-side had one ribbon, whereas 7 had double ribbons. In contrast to GluA3^WT^, 3 synapses on the pillar-side in GluA3^KO^ mice had 2 ribbons, whereas 8 had a single ribbon. We did not find differences in PSD surface area or volume between the modiolar or pillar synapses with one or two ribbons, so data were combined. We then compared the PSD surface area and volume between the modiolar and pillar sides. All statistical tests in Fig 8 are unpaired t-test unless noted. Similar to the GluA3^WT^, there was no significant difference in the PSD surface area (p = 0.16) or volume (p = 0.31) (**Fig. 8A bottom; Table 1**). Unlike the GluA3^WT^, the PSD length was significantly greater on the modiolar than on the pillar side (p = 0.028, Mann-Whitney test) (**Fig. 8B bottom; Table 2**).

The analysis of presynaptic ribbon surface area showed that those on the modiolar side had significantly larger surface area than those on the pillar side (p = 0.007) (**Fig. 8A bottom; Table 1**). In contrast, ribbon volume was similar between the modiolar and pillar sides (p = 0.12) (**Fig. 8A bottom; Table 1**).

Like GluA3^WT^ females, the major axis of the KO, was similar between the modiolar- and pillar-side ribbons (p = 0.08) (**Fig. 8B bottom; Table 2**). In contrast to GluA3^WT^ mice, the minor axis of the KO, was similar between the ribbons of the modiolar- and pillar-side synapses (p = 0.15) (**Fig. 8B bottom; Table 2**). Pillar-side ribbons were more circular (p = 0.001) and round (p = 0.005) than modiolar-side ribbons (**Fig. 8B bottom; Table 2**), but the difference was less significant than in the GluA3^WT^. The analysis of the SVs size on modiolar- and pillar-side synapses showed that they were similar (p = 0.63, Mann-Whitney test) (**Fig. 8B**; **Table 2**).

### SV size and ribbon shape were altered in the 5-week-old female GluA3^KO^ mice in quiet

Next, we compared the pre- and post-synaptic ultrastructural features on the modiolar- and pillar-side synapses between genotypes (unpaired t-test in all the comparisons). The PSD surface area and volume were similar on the modiolar (surface area p = 0.43; volume p = 0.16) and pillar side synapses (surface area p = 0.78; volume p = 0.74) between GluA3^KO^ and GluA3^WT^ mice (**Fig. 9A left**). No differences in PSD length were observed between genotypes (modiolar p = 0.68; pillar p = 0.11) (**Fig. 9B top left**).

**Figure 9.**
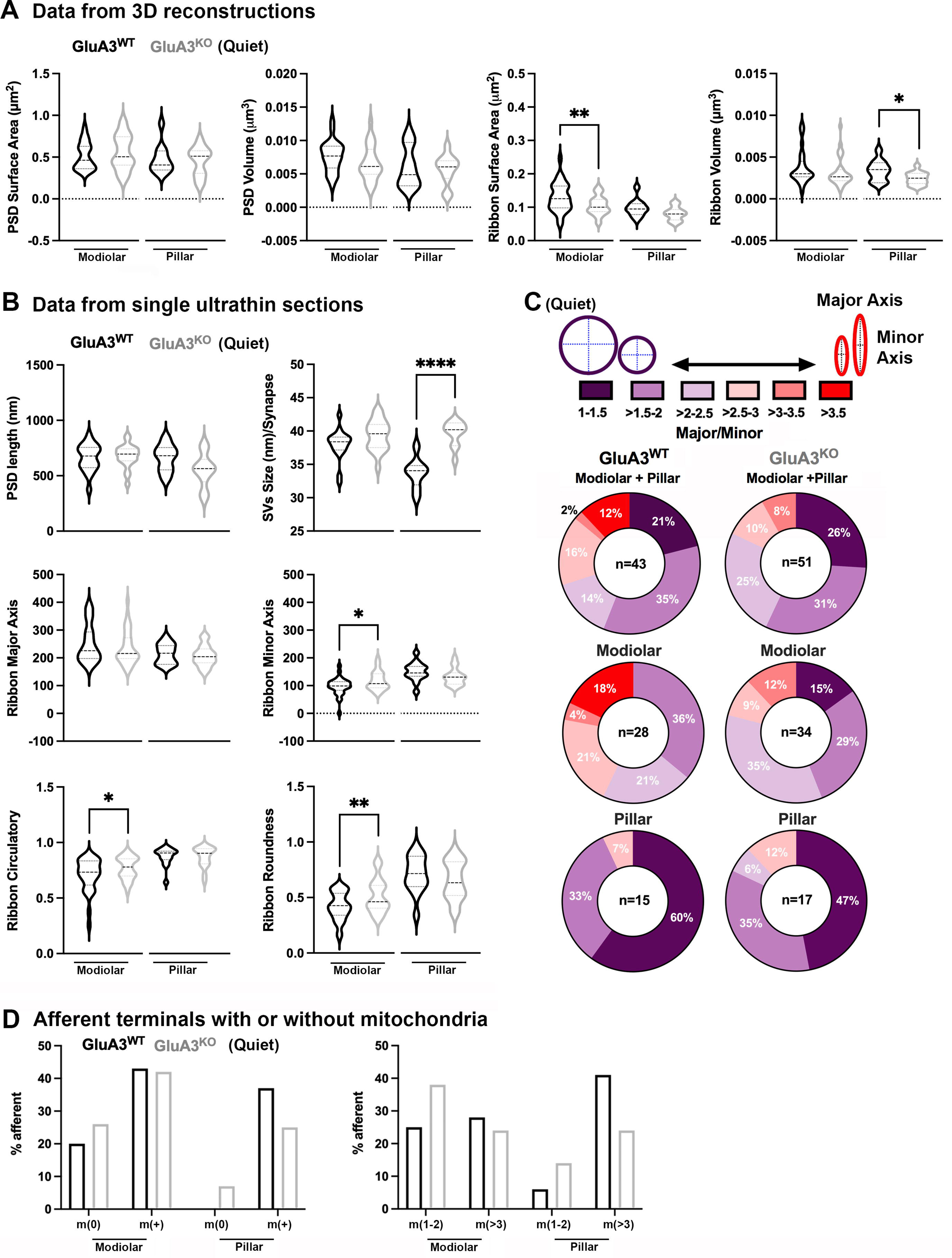
SV size and pre-synaptic ribbon shape are altered in the 5-week-old GluA3^KO^ female mice in quiet. **A)** Violin plots show the quantitative comparison of the surface and volume of PSDs and ribbons obtained from 3D reconstruction between GluA3^WT^ (black) and GluA3^KO^ (gray) mice. One-way ANOVA, *p < 0.05, Data corresponds to median. **B)** Violin plots show the quantitative comparison of PSD length, major and minor ribbon, ribbon circularity and roundness, and synaptic vesicle (SV) size per synapse between GluA3^WT^ (black) and GluA3^KO^ (gray). One-way ANOVA, *p < 0.05, **p < 0.01, ***p < 0.001, ****p < 0.0001, Data corresponds to median. **C)** Pie charts show the variability in ribbon shape in percentage from round (red) to elongated (purple) as a total (modiolar + pillar), and then separated by modiolar and pillar-side ribbons of GluA3^WT^ (n = 43) and GluA3^KO^ (n = 51). The Major Axis / Minor axis ratio denotes the ribbon shape regardless of the size, as explained in a schematic drawing on the top of the figure. **D)** Histograms show the percentage of modiolar and pillar side afferent terminals without and with mitochondria (m (0) and m+, respectively) in the GluA3^WT^ (n=32) and GluA3^KO^ (n=29) mice raised in quiet (left). The histogram on the right shows the percentage of modiolar and pillar side afferent terminals with 1-2 and more than 3 mitochondria per terminal.

Synaptic ribbon surface area on the modiolar side was significantly reduced in GluA3^KO^ compared to GluA3^WT^ (p = 0.0096). In comparison, the surface area on the pillar side was similar between the genotypes (p = 0.073) (**Fig. 9A right**). Ribbon volumes on the modiolar side were similar between GluA3^KO^ and GluA3^WT^ (p = 0.25), whereas, on the pillar side, the ribbon volumes of GluA3^KO^ were significantly smaller than GluA3^WT^ (p = 0.029) (**Fig. 9A right**). There was no significant difference in ribbon major axes on the modiolar-or pillar-side synapses between the GluA3^WT^ and GluA3^KO^ (modiolar: p = 0.34, pillar: p = 0.56). In contrast, the ribbon minor axes of GluA3^KO^ modiolar side synapses were larger than those of the GluA3^WT^ (p = 0.018). The minor axes of pillar side synapses were the same between genotypes (p = 0.12) (**Fig. 9B center**). The ribbon of modiolar side synapses of GluA3^KO^ was more circular and rounder than in GluA3^WT^ (circularity, p = 0.049, roundness, p = 0.010). On pillar-side synapses, ribbon circularity and roundness were similar between the genotypes (circularity, p = 0.90, roundness, p = 0.36) (**Fig. 9B bottom**). While SV size on the modiolar side was similar between genotypes, those on the pillar side were significantly larger in GluA3^KO^ mice (modiolar, p = 0.059, pillar, p < 0.0001) (**Fig. 9B top right**).

Next, we designed an analysis in 2D to determine the ribbon shape by dividing the major to the minor axis (see Material and Methods); this ratio is independent of size. The schematic diagram in Figure 9C explains that round ribbons (shown in dark purple) have equal lengths on the major and minor axes. In contrast, elliptical ribbons (shown in red) have a major axis longer than the minor. The pie charts show the variability in ribbon shape from round to elongated as a total (modiolar + pillar) and then separated by modiolar- and pillar-side ribbons (**Fig. 9C**). Overall, ribbons in GluA3^WT^ and GluA3^KO^ varied in shape from round to elongated; however, the KO ribbons were less elongated and rounder (**Fig. 9C top pie chart**). When comparing the shapes separated by modiolar and pillar, data showed that 18% of the modiolar-side ribbons in the GluA3^WT^ mice were elliptical, although a larger percentage (36%) were round. In the WT pillar-side ribbons, the large majority (60%) were round, and the elliptical ribbons were not observed (**Fig. 9C**). In the GluA3^KO^, in contrast to the GluA3^WT^, we did not find the most elliptical category within the modiolar-side ribbons. In the KO, 15% of modiolar ribbons were round, and the rest (85%) were between elliptical and round. Almost half (47%) of pillar-side ribbons in the GluA3^KO^ were round, and the others had intermediary shapes (**Fig. 9C**).

In summary, our ultrastructural data from 5-week-old female mice raised in quiet showed that in contrast to the GluA3^WT^, the PSD length of the GluA3^KO^ modiolar and pillar side synapses differed (smaller in the pillar). However, this difference was not significant when compared between genotypes. In addition, we observed that the shape of the pre-synaptic ribbons of both synapse types and the SV size of pillar-side synapses were altered in the female GluA3^KO^ mice in quiet.

### Type I afferent terminals in the GluA3^KO^ mice in quiet had a decrease in mitochondria content

Mitochondria often occupy neurons and have many protective properties, including their role in calcium buffering (Duchen, 2000; Wang et al., 2019; Matuz-Mares et al., 2022; Verma et al., 2022). To analyze the presence of mitochondria in the afferent terminals of Type I SGNs on IHCs, we quantified the number and percentage of mitochondria in modiolar- and pillar-side afferents of GluA3^WT^ and GluA3^KO^ mice in quiet (**Fig. 9D left**). In GluA3^WT^, out of the total afferent terminals analyzed along modiolar and pillar side of hair cells, 20% of modiolar-side afferent terminals were without mitochondria, whereas 43% had (range: 1 - 9 mitochondria per afferent; median: 3). In contrast, all the pillar-side afferent terminals (representing 37% of the total) had mitochondria (range: 1 - 8 mitochondria per afferent; median: 4). The analysis of the GluA3^KO^ shows that out of the total, 26% of the modiolar-side afferents lacked mitochondria, whereas most had them (42% of the total; range: 1 - 8 mitochondria per afferent; median: 2). Of the pillar-side afferents, we found that in contrast to the GluA3^WT^, 7% lacked mitochondria whereas 25% had them (range: 1 - 8 mitochondria per afferent; median: 4). Overall, we find that in GluA3^KO^ mice, there is an increase in the percentage of both modiolar- and pillar-side afferents lacking mitochondria compared to GluA3^WT^ mice.

Within the afferents with mitochondria, either in GluA3^WT^ or GluA3^KO^ mice, the number of mitochondria varied from 1 to 9. Because the median was around 3 on average, we determined how many afferents had fewer mitochondria below the median (n = 1 - 2 mitochondria per afferent) and how many had above the median (more than 3) (**Fig. 9D right**). In GluA3^WT^, the percentage of modiolar-side afferent terminals with a lower and higher number of mitochondria was similar (25% and 28%, respectively). In contrast, only a small percentage (6%) of pillar-side afferent terminals had a lower number of mitochondria, whereas the majority (41%) had a higher number. In the GluA3^KO^, the percentage of modiolar-side afferents with one or two mitochondria was larger than those with more than three (38% and 24%, respectively). In pillar-side afferents, 14% had one or two mitochondria, while the rest (24%) had more than three. When comparing GluA3^WT^ and GluA3^KO^ data, we found that in the GluA3^KO,^ there was an increase in the number of modiolar and pillar side afferents with a low number of mitochondria. A larger percentage of afferent terminals without or with a lower number of mitochondria in the female GluA3^KO^ could contribute to a higher IHC-ribbon synapse vulnerability to ambient sound levels.

## Discussion

In the present study, we analyzed the role of the GluA3 subunit on sound encoding, AMPAR subunit composition, and pre-and post-synaptic ultrastructure of IHC-ribbon synapses in female mice in ambient and quiet sound conditions. Our study shows that in the female GluA3^KO^ mice, only ambient background sound levels lead to early hearing impairment (**Figs. 1 and 2**), an increase in the number of ribbonless synapses (**Fig. 3**), and afferent terminal swellings (**Fig. 7; Fig. EV2**), suggesting excitotoxicity. The excitotoxicity in afferent terminals of female GluA3^KO^ potentially results from an increase in the Ca^2+^ permeability through the AMPAR channel as a consequence of the higher GluA4:GluA2 ratio and altered intrasynaptic distribution of the subunits within the postsynaptic structure of IHC-ribbon synapses (**Figs. 4, 5 and 6; Fig. EV1**). In addition, we find that GluA3 plays a crucial role in maintaining the morphology of pre-synaptic ribbons and SVs, and of afferent terminal mitochondria content in both modiolar- and pillar-side synapses (**Figs. 8 and 9**), all indicating that GluA3 is necessary for the development and maturation of IHC-ribbon synapses. Together, our results indicate that the AMPAR subunit GluA3 is required to prevent excitotoxicity in ambient sound levels and maintain the normal structure of IHC synapses in young adult female mice.

### Lack of GluA3 in females increases IHC-ribbon synapse vulnerability to ambient sound

We recently found that 5-week-old GluA3^KO^ males raised in ambient sound levels have normal hearing sensitivity, measured by ABR click and pure-tone thresholds and wave-1 amplitude (Rutherford et al., 2023). The male GluA3^KO^ mice do not show hearing impairment until they reach 2 months of age (García-Hernández et al., 2017). Here, we show that female GluA3^KO^ raised in the same ambient background sound levels have a decreased wave-1 amplitude at 5 weeks of age that persists up to at least 3 months (**Fig. 1**). This early decrease in wave-1 amplitude suggests reduced SGN synchrony in the female GluA3^KO^ mice. Interestingly, our data also show that the increased vulnerability of the female GluA3^KO^ to ambient sound can be prevented at least up to 3 months of age if mice are raised in much quieter sound environments (**Figs. 1 and 2**). All of this suggests that female GluA3^KO^ mice show a greater vulnerability than males to changes in sound environment.

Among the potential contributors to increased hearing loss vulnerability in the absence of GluA3 is a change in AMPAR subunit composition, specifically less GluA2 or more GluA4 (**Figs. 3 - 6**), likely to increase the abundance of Ca^+2^-permeable AMPARs (CP-AMPARs) at the IHC-ribbon synapse. AMPARs underlie fast excitatory synaptic transmission at IHC-ribbon synapses (Ruel et al., 1999: Glowatzki and Fuchs, 2002). The activity of AMPARs is crucial for the normal physiology of synapses and, on the other hand, for the induction of excitotoxicity (Puel et al., 1998; Ruel et al., 2000). Several studies have shown that noise-induced hearing loss can produce excitotoxicity through excessive release of glutamate and subsequent activation of AMPARs, specifically Ca^2+^-permeable AMPARs (Kim et al., 2019; Hu et al., 2020). AMPARs at mature IHC-ribbon synapses are assembled as tetramers comprised of different combinations of GluA2 – 4 subunits; homotetramers can exist but are less stable energetically (Rossmann et al., 2011; Zhao et al., 2017). Calcium conductance of AMPARs varies depending on the presence of GluA2 subunits in the tetrameric complex, and AMPARs lacking GluA2 subunits are Ca^+2^ permeable (Hollmann et al., 1991; Geiger et al., 1995). Immunoreactivity for GluA2/3 or GluA2 is widely reported as present at all, or nearly all, IHC-ribbon synapses (Khimich et al., 2005; Meyer et al., 2009; Liberman et al., 2011). The presence of GluA2 presumably reduces overall synaptic Ca^2+^ influx and vulnerability to glutamate excitotoxicity. Nonetheless, glutamate excitotoxicity underlies noise-induced cochlear synaptopathy, and recent studies in rats and mice have shown the existence of apparently GluA2-lacking nanodomains presumably containing CP-AMPARs within postsynaptic densities and that antagonizing these CP-AMPARs can prevent noise-induced synaptopathy (Sebe et al., 2017; Hu et al., 2020).

While the full complement of AMPAR pore-forming and auxiliary subunits functioning at IHC-ribbon synapses is still undetermined, in the absence of GluA3 and GluA1 (Rutherford et al., 2023), GluA2/4, GluA2/2, and GluA4/4 are the only possible pore-forming dimers to dimerize into the tetrameric AMPAR complex. In our study, we find the number of paired synapses (i.e., a presynaptic ribbon paired with postsynaptic GluA2 and GluA4) was similar between GluA3^WT^ and GluA3^KO^ female mice in both ambient and quiet rearing by 5 weeks (**Fig. 3**). However, the number of lone ribbons and ribbonless synapses was significantly greater in the KO, but interestingly, only if raised in ambient sound levels (**Fig. 3**). GluA2-lacking synapses were not observed in GluA3^WT^ and in less than 2% of synapses in GluA3^KO^ in quiet or ambient. GluA4-lacking synapses (i.e., a presynaptic ribbon paired with postsynaptic GluA2 only) were not observed in quiet in either genotype. Ambient sound increased the occurrence of GluA4-lacking synapses in both genotypes, with significantly more in GluA3^WT^ than GluA3^KO^ (∼ 30% of synapses in some images from the 10 kHz region (**Fig. 3**). Thus, ambient sound appears to induce a shift toward GluA2-only synapses in GluA3^WT^ and toward unpaired pre- and post-synaptic puncta in GluA3^KO^. Both genotypes had unpaired synapses when reared in ambient sound, but ribbonless synapses containing GluA2 and GluA4 were more abundant in GluA3^KO^ (∼ 10%) than GluA3^WT^ (∼ 5%). As well, these ribbonless synapses in GluA3^WT^ tended to have greater GluA4:GluA2 ratios (implied Ca^2+^ permeability) than the paired synapses in the same images but not in GluA3^KO^ (**Fig. 4**). If the presence of the ribbon in GluA3^WT^ is associated with AMPARs having relatively less Ca^2+^ permeability, one implication is that ambient sound recruits GluA2 to ribbon synapses, somehow making them relatively less vulnerable to excitotoxicity.

In GluA3^KO^ females, as in males, the ribbon-occupied synapses (i.e., paired synapses) are already elevated in the GluA4:GluA2 ratio, suggesting an increase in the relative abundance of GluA2-lacking CP-AMPARs (**Fig. 4**). This may lead to excessive influx of Na^+^ and Ca^2+^ into the Type I afferent terminals, increasing their intracellular concentration. Intriguingly, the increase in the GluA4:GluA2 ratio occurred in ambient and in quiet conditions, but the change was larger in ambient (∼ 103% increase in ambient vs. ∼ 56% increase in quiet). Remarkably, only in the female KO in ambient conditions did we observe afferent swellings (**Fig. 7; Fig. EV2**), which were similar to but much less extensive than those reported after noise exposure or following cochlear perfusion with excitotoxic glutamate receptor agonists (Robertson, 1983; Puel et al., 1998). Thus, the further increase in the GluA4:GluA2 ratio between quiet and ambient seems sufficient to trigger a slowly progressing excitotoxicity in the afferent terminals. Structural features of swollen afferents include missing mitochondria, swollen endoplasmic reticulum, and decreased cytoplasm electron-density. Some swellings had a PSD in opposition to a pre-synaptic ribbon, while others lacked the ribbon. The former would likely appear as a normal paired synapse in confocal imaging, and the latter would be considered a ribbonless synapse. Altogether, our results show the existence of chronic ultra-structural and physiological ambient sound-induced synaptopathy long before the synapses degenerate in the female GluA3^KO^ cochlea.

### Molecular and ultrastructural changes in GluA3^KO^ females are already present in quiet

Even in the absence of ambient sound, we found differences between GluA3^WT^ and GluA3^KO^. In addition to the greater overall GluA4:GluA2 fluorescence ratios in GluA3^KO^ synapses, we find an altered intrasynaptic distribution of ratios (females in quiet, **Figs. 5 - 6**). In GluA3^WT^, the ratios tended to peak 200 – 400 nm from the synapse center, which would be closer to the perimeter of the PSD as measured in TEM (**Fig. 7**). This peak was less pronounced in GluA3^KO^ because ratios tended to be greater near the synapse center in GluA3^KO^ than in GluA3^WT^. Electron tomography has localized SVs morphologically docked to the presynaptic membrane only within ∼ 100 nm of where the synaptic ribbon is anchored to the active zone, suggesting exocytosis may be limited to within 100 nm from the ribbon (Chakrabarti et al., 2022). More GluA2-lacking and presumably CP-AMPARs residing closer to the ribbon would likely increase the probability of their activation after presynaptic glutamate release, perhaps leading to excitotoxicity in female GluA3^KO^ mice raised in ambient sound levels.

Postsynaptic calcium homeostasis is crucial to protect synapses from excitotoxic insult (Verma et al., 2022), for which mitochondria play an important role (Duchen, 2000; Verma et al., 2022). Reduced calcium buffering by mitochondria in the afferent terminals of GluA3^KO^ females could contribute to afferent terminal swellings and excitotoxicity. Structural studies in cats and mice indicated that pillar-side terminals contained more and larger mitochondria than modiolar-side terminals (Liberman, 1980; Moverman et al., 2023). This difference in mitochondrial content might increase the vulnerability of afferent terminals on the modiolar side to damage after noise exposure. In our study, we find a similar trend in the female C57BL/6 GluA3^WT^ mice raised in quiet, where pillar-side terminals contained more mitochondria than modiolar-side terminals (**Fig. 9D**) In addition, we identified a ∼20% of modiolar afferents without mitochondria at the afferent terminals in the GluA3^WT^. However, in the GluA3^KO^ females, on both modiolar- and pillar-side synapses we find an increase in afferent terminals either without mitochondria (within the image field of view) or with a lower number of mitochondria (**Fig. 9D**). The afferent terminals of female GluA3^KO^ raised in quiet are structurally and functionally asymptomatic (i.e., no swellings, no ABR threshold or wave-1 phenotype), but these results suggest less Ca^2+^ buffering capacity, which may increase vulnerability in ambient sound. Postsynaptic mitochondria are critical to the development, plasticity, and maintenance of synaptic inputs (Thomas et al., 2023). Although the link between reduced mitochondrial content and the absence of GluA3 is unclear, our results suggest GluA3 may influence modiolar- and pillar-side differences in mitochondrial content that emerge during development. Altogether, the altered synaptic AMPAR subunit composition and the decrease in mitochondria content at afferent terminals of the female GluA3^KO^ in quiet, while being asymptomatic, could increase the synapse vulnerability if mice are raised in louder ambient sound levels, as we find in this study.

### Lack of modiolar-pillar spatial gradient of synaptic structural heterogeneity at IHC-ribbon synapses in GluA3^KO^ mice

Our recent ultrastructural and imaging study of cochlear ribbon synapses in 5-week-old male GluA3^KO^ mice included dysregulation of GluA2 and GluA4 subunit relative abundance and alterations in pre- and post-synaptic ultrastructure associated with an increased vulnerability to glutamatergic synaptopathy at ambient, background levels of sound (Rutherford et al., 2023). These structural and molecular alterations at the cochlear ribbon synapses of pre-symptomatic 5-week-old male GluA3^KO^ mice appear to be pathological, preceding the reduction in ABR wave-1 amplitudes observed at 2-months of age (Garcia-Hernández et al., 2017). Among the structural alterations, we found that IHC modiolar-pillar differences (larger ribbons on the modiolar side than the pillar side) were eliminated or reversed in the male GluA3^KO^. Specifically, the major axis of the modiolar-side ribbons was smaller in GluA3^KO^ males. The SVs were also larger on both the modiolar and pillar sides compared to the GluA3^WT^ (Rutherford et al., 2023). Here, our structural analysis of 5-week-old female GluA3^WT^ and GluA3^KO^ in ambient and quiet showed that only the female KOs raised in ambient sound levels had decreased wave-1 amplitude and the characteristic Type I terminal swellings, indicating functional and structural synaptopathy (**Figs. 1, 2 and 7**). Thus, we reasoned that analyzing the structure of the IHC-ribbon synapses of the female KOs in quiet would provide a unique opportunity to identify pre-symptomatic features associated with an increased vulnerability to glutamatergic synaptopathy at ambient sound levels.

Our ultrastructural analysis of the IHC-ribbon synapses of females in quiet showed that in C57BL/6 GluA3^WT^ mice, the PSD and ribbon major axis lengths of the modiolar and pillar side synapses were similar (**Figs. 8 and 9**), differing from what we and others found in WT C57BL/6 or CBA/CaJ males (Rutherford et al., 2023; Liberman et al., 2011) or in a (presumably male) cat raised in a low-noise chamber (Liberman, 1980), where modiolar side ribbons are larger. In GluA3^KO^ females, the modiolar-side ribbons widened due to a significant increase in the minor axis length, while pillar ribbons tended to be more elongated due to the reduced minor axis length, and no difference was observed in the ribbon major axis (**Figs. 8 and 9**). Similar to the male GluA3^WT^, the SVs were larger on the modiolar side in female GluA3^WT^. In GluA3^KO^, the SVs on modiolar and pilar side synapses did not differ in size because SVs on the pillar side were larger. The length of the PSD in female GluA3^KO^ mice was larger in modiolar-side synapses compared to pillar side (**Figs. 8 and 9**). Altogether, we find that lack of GluA3 alters pre- and postsynaptic features of both modiolar- and pillar-side synapses in both sexes. However, we were surprised that the effects in the female GluA3^KO^ were less prominent than in the males, especially the structural alterations of the modiolar side ribbons. This could be because we analyzed the females reared in the quiet, while our previous study assessed male mice raised in ambient sound. In any case, the spatial gradient of modiolar- and pillar-side synapse heterogeneity in WT seems weaker in females than males. Nonetheless, our ultrastructural analysis in females supports the role of GluA3 in maintaining pre- and post-synaptic structural features. We propose that this occurs through an unknown trans-synaptic mechanism during development at IHC-ribbon synapses in the cochlea and the endbulb of Held synapses in the cochlear nucleus (García-Hernández et al., 2017; Antunes et al., 2020; Rutherford et al., 2023). AMPAR subunits in the cochlea may interact with the trans-synaptic adhesion proteins Neuroligins and Neurexins, which could also interact with presynaptic proteins and voltage-gated Ca^2+^ channels at IHC-ribbon synapses (Arac et al., 2007; Luo et al., 2020; Ramirez et al., 2022).

### Why are females lacking GluA3 more vulnerable to ambient sound levels than males?

Studies in mice, chinchillas, and humans reported that males are more vulnerable to noise-induced hearing loss than females (McFadden et al., 2000; Milon et al., 2018; Szanto and Ionescu, 1983; Shuster et al., 2021). Here, we show that the absence of the GluA3 AMPAR subunit increases the vulnerability of the female mice to ambient sound levels. Taken together, our data suggests that the decreased vulnerability observed in female mice requires the presence of GluA3 at the IHC-ribbon synapse. The effect of GluA3’s absence on increasing excitotoxicity in female mice reared in ambient sound may be indirect due to the decrease in GluA2 at the synapse **Figs. 3 - 6**); however, this decrease in GluA2 was observed also in male GluA3^KO^ mice reared in ambient sound (Rutherford et al., 2023). Perhaps females have a unique requirement for GluA3, or males have a unique means of resisting excitotoxicity in its absence. GluA3 and GluA2 form the most predominant dimers in the brain (Rossmann et al., 2011; Zhao et al., 2017), and we assume this biophysical property extends to the inner ear, although this remains to be shown directly. We speculate that loss of GluA3 results in decreased trafficking of GluA2 to the synapse since *Gria2* RNA levels were not reduced in *Gria3^KO^*mice (Rutherford et al., 2023). Still, it remains unclear why the 5-week-old GluA3^KO^ male mice did not exhibit a reduction in ABR wave-1 amplitudes or elevation of ABR thresholds. Recently, we showed that spiral ganglion neurons from female mice have a two-fold higher expression of *Gria3* mRNA and larger ABR wave-1 amplitudes than male mice (Lozier et al., 2023). While *Gria2* and *Gria4* are located on autosomes, *Gria3* is on the X chromosome (Mahadevaiah et al., 2009). Therefore, it is possible that females have more GluA3 due to incomplete X-inactivation in some or all SGNs, leading to increased gene dosage (Carrel and Willard, 2005; Berletch et al., 2015). If so, knocking out *Gria3* could have a greater impact on the female than the male IHC-ribbon synapse.

## Material and Methods

### Animals

In this study, a total of 57 female C57BL/6 wild type (GluA3^WT^, n= 26) and GluA3-knockout (GluA3^KO^, n= 31) mice from two different colonies were used between postnatal days 20 (P20, 3-weeks) to 90 (P90, 13-weeks). The generation of the GluA3^KO^ mice has been previously described (García-Hernández et al., 2017; Rubio et al., 2017). Mice were raised up to P20 in quiet sound pressure levels conditions (40 - 55 dB SPL, 100 Hz – 8.3 kHz, general audible range for humans, measured with an audiometer, General Tools & Instruments LLC; 10 dB SPL, 3 kHz–90 kHz, mouse hearing range, Sensory Sentinel, Turner Scientific, Jacksonville, IL). One group of GluA3^WT^ (n= 13) and *GluA3*^KO^ (n= 14) mice were maintained in the same quiet conditions until P90 (**Fig. 1A**). A second group of mice (GluA3^WT^ n= 13) and (GluA3^KO^ n= 17) were moved at P20 to a room facility with ambient sound pressure levels (55-75 dB SPL,100 Hz – 8.3 kHz, general audible range for humans measured with an audiometer; 40 dB SPL, 3 kHz– 90 kHz, mouse hearing range, Sensory Sentinel) and maintained until they reached P90 (**Fig. 1A**). Both mouse groups were reared in a 12-hour light/12-hour dark daily photoperiod and were fed *ad libitum*. Auditory brainstem recordings (ABRs) were collected at four postnatal days: P20, P35, P60, and P90. Cochleae were dissected from P35 mice and used for further experiments (**Fig. 1A**). All experimental procedures were per the National Institute of Health guidelines and approved by the University of Pittsburgh Institutional Animal Care and Use Committee.

### Auditory Brainstem Recordings (ABRs)

At least 10-15 GluA3^WT^ and GluA3^KO^ mice from each animal group (quiet and ambient) and age group were used for ABRs. During the ABRs, mice were kept anesthetized with isoflurane (3% induction and 1.5% maintenance), and the internal temperature was monitored with a thermometer (J/K/T Thermocouple, Kent Scientific Corporation) that was kept constant between 37-38 ^°^C using isothermal heat pads. Open-field ABRs and analysis were performed as described in Lozier et al. (2023). Briefly, ABRs were recorded in a sound attenuation chamber inside a Faraday cage. The speaker was calibrated in the chamber using a microphone (PCB electronics, model no. 377C01, Depew, NY) placed 10 cm from the multi-field magnetic speaker (MF1, Tucker-Davis Technologies (TDT), Alachua, FL), which was the same distance as the mouse’s external ear to the speaker during experiments. For ABR recordings, needle electrodes were placed subcutaneously at the vertex of the scalp and below the pinna of both the tested (ipsilateral) and contralateral ears. Clicks (1 ms) or pure tones of 4, 8, 12, 16, 24, and 32 kHz (0.5 ms) stimuli were presented from 90 dB to 10 dB in decreasing steps of 5 dB at 21 sweeps with an interstimulus interval of 47.6 ms. All the recordings’ responses were averaged over 512 sweeps, amplified, digitized, and transferred via an optical port to the RZ6 processor. ABR click, and pure tone thresholds were determined as the lowest intensity level (in dB SPL) at which reproducible peaks were visible. The clicks wave I amplitude (in μV) was determined as the difference between the peak and subsequent trough.

### Immunofluorescence

A total of 9 female mice (GluA3^WT^ n = 4; GluA3^KO^ n = 5) at 5 weeks of age, raised in either ambient sound (ambient group) or low-level sound (quiet group) were anesthetized with ketamine (60 mg/kg) and xylazine (6.5 mg/kg), then perfused intravascularly for 10 minutes with fixative consisting of 4% paraformaldehyde (PFA) in 0.1M phosphate buffer (PB), pH = 7.2. Temporal bones were then isolated from the skull, the stapes was removed, the oval and round window membranes were punctured, and a hole was opened at the apex of the cochlear bone near the helicotrema. Each cochlea was perfused with fixative through the oval window while bathing in a dish of fixative. After postfixation for 45 minutes in the dish on top of an ice pack, the cochleae were rinsed in 0.1M phosphate-buffered saline (PBS) and then shipped to Washington University in St. Louis overnight in PB supplemented with 5% glycerol.

GluA3^WT^ and GluA3^KO^ cochleae were batch processed in parallel using the same reagent solutions in two separate groups, one raised in ambient background sound and the other raised in quiet background sound levels. After rinsing in PB, each temporal bone was decalcified in 50 mL of PB supplemented with 10% EDTA for 2 hours at room temperature on a rocker. After removing the outer shell of the cochlea and peeling away the lateral wall, the spiral containing the organ of Corti, osseous spiral lamina, and spiral ganglion was isolated and cut into apical-cochlear, middle-cochlear, and basal-cochlear pieces for wholemount immunolabeling as previously described (Jing et al., 2013; Ohn et al., 2016; Sebe et al., 2017; Kim et al., 2019; Hu et al., 2020; Rutherford et al., 2023). To permeabilize the tissue and to block non-specific binding of antibodies, the samples were immersed overnight at 4°C in 100 µL of blocking buffer (0.3% Triton X-100 and 16% normal horse serum in PBS) in a 9-well Pyrex glass plate. To label presynaptic ribbons (CtBP2/Ribeye) and postsynaptic AMPA-type glutamate receptor subunits (GluA2, GluA3, and GluA4), primary antibodies were diluted in blocking buffer (1:400) and samples were incubated in the 9-well plates with 100 μL per plate overnight at 4°C: CtBP2 mouse IgG1 (BD Biosciences 612044; RRID:AB_399431), GluA2 mouse IgG2a (Millipore MAB397; RRID:AB_2113875), GluA3 goat (Santa Cruz Biotechnology SC7612), and GluA4 rabbit (Millipore AB1508; RRID:AB_90711). After rinsing 3 times in wash buffer (PBS supplemented with 0.3% Triton-X), samples were incubated for 1 hour at room temperature in species-appropriate secondary antibodies diluted in blocking buffer (1:100). Secondary antibodies were pre-conjugated to Alexa Fluor (Life Tech.) fluorophores excited by 488, 555, or 647 nm light. After rinsing 3 times in wash buffer and PBS, the 3 pieces from each cochlea were mounted in Mowöl between a microscope slide and coverslips (#1.5, Zeiss). Slides were kept at 4°C until imaging.

### Conventional Confocal Microscopy and Image Analysis

Confocal volumes were acquired with 12-bit images, 100 µm wide containing approximately 12 IHCs, on a Zeiss LSM 700 with a 63X 1.4 NA oil objective lens with a Z-step of 0.37 µm and pixel size of 50 nm in X and Y, and a pinhole setting of 1 AU. We first surveyed the samples to determine the necessary laser power settings to collect all the images at a gain of 700 while avoiding pixel intensity saturation. Using identical acquisition settings across samples, we collected 3–4 images from the center of each of the 3 pieces (basal-, middle-, and apical-cochlear) near tonotopic characteristic frequencies of 10, 20, and 40 kHz (Müller et al., 2005). For display only, the brightness and contrast levels were adjusted linearly for visual clarity. Image analysis was performed on raw data.

The numbers of hair cells and synapses were counted and manually verified after semi-automated identification using Imaris software (Bitplane) to calculate the mean number of synapses per IHC per image. The observers were blinded to mouse genotype. For each group of images, group means (± SD) were calculated across image means. Paired synapses were identified as juxtaposed puncta of presynaptic ribbons (Ribeye/CtBP2) and postsynaptic AMPA receptor patches (GluA2 and/or GluA4), which appear to partly overlap at confocal resolution (Rutherford, 2015). Unpaired (i.e., lone) ribbons were defined as Ribyeye/CtBP2 puncta located under inner hair cell nuclei, in the synaptic region, but lacking apposed GluA2 or GluA4 puncta. Inner hair cell nuclei were labelled with CtBP2/Ribeye because CtBP2 is a transcriptional co-repressor. For unpaired ribbons, we did not distinguish membrane-anchored from unanchored. Ribbonless synapses consisted of GluA2 and/or GluA4 puncta located in the synaptic region under the inner hair cell nuclei but lacking apposed presynaptic ribbons.

Pixels comprising puncta of synaptic fluorescence were segmented in 3D as ‘surface’ objects in Imaris using identical settings across image stacks, including the ‘local contrast background subtraction’ setting to automatically calculate the appropriate pixel-intensity threshold for each fluorescence channel in each image. This adaptive and automatically-calculated thresholding algorithm compensated for differences in overall luminance between image stacks that would affect the volume of segmented puncta if a fixed threshold was applied across images, and avoided the potential subjective bias of setting a user-defined arbitrary threshold value separately for each image. Intensity per synaptic punctum was calculated as the summation of pixel intensities within the surface object. To associate the intensities of puncta belonging to the same synapse, we generated surface objects from a 4^th^ channel calculated in Imaris as the sum of the three channels (GluA2, 3, and 4; or CtBP2, GluA2, and GluA4). Each of these 4^th^-channel objects contained the entire synapse (presynaptic and postsynaptic elements). The summated pixel intensities for each of the three fluorescence channels were calculated within each synapse. Some synapses appeared to lack a presynaptic ribbon. The CtBP2 channel was used to make surface objects of ribbons. These ribbon objects were used to remove ribbonless synapses from the 4th-channel objects by eliminating 4^th^-channel objects with centers further than 0.1 µm from the center of the nearest ribbon. The average density of synaptic fluorescence per image was calculated as median punctum intensity (a.u.) divided by median punctum volume (µm^3^) using surface objects calculated from corresponding individual fluorescence channels.

### Airyscan Confocal Microscopy and Image Analysis

Airyscan volumes were acquired with 16-bit images, 46 µm wide containing approximately 5 IHCs, on a Zeiss LSM 880 Confocal with Airyscan, with a 100X 1.5 NA oil objective lens with a Z-step of 0.168 µm and pixel size of 26 nm in X and Y, and a pinhole setting of 1.5 AU. We collected stacks near tonotopic characteristic frequencies of 11, 20, 25, 38, 57, and 70 kHz using identical acquisition settings for all images. For display only, the brightness and contrast levels were adjusted linearly for visual clarity. Image analysis was performed on unadjusted data after Airyscan Processing.

To quantify changes in the relative abundance of GluA2 and GluA4 around individual synapses, 3-dimensional data was reduced to 1-dimensional normalized radial distributions of fluorescence density ratios around the synapse center. Analysis was performed blinded on the maximum intensity Z-projections for spatially-isolated synapses only (i.e., synapses having no overlap with other synapses XY or Z). The 2-dimensional centroid of the synaptic ribbon was defined as the synapse center. Radial distributions of normalized GluA4: GluA2 fluorescence density ratio as a function of distance from the synapse center were constructed with custom code in MATLAB Software. For each fluorescence channel, the sum of pixel intensities was calculated for a 3 × 3 pixel box (78 × 78 nm) centered on the ribbon centroid and then for 11 shells around the center box, with each shell expanding by 2 pixels (52 nm) on each side, such that the first shell was outlined by a square with dimensions of 7×7 pixels (182 × 182 nm) and the largest shell was outlined by a square of 47×47 pixels (1,222 × 1,222 nm). For each shell, the area was calculated as the number of pixels, and the sum of pixel intensities was divided by the area to calculate the ‘shell density’ (intensity / area) for each fluorescence channel. For each isolated synapse in an image, the GluA4: GluA2 radial distribution of the ‘ratio of raw shell densities’ was calculated as the blue shell density divided by the green shell density. The peak-normalized shell densities for each synapse were calculated by normalizing to the maximum shell density of each fluorescence channel such that each radial distribution of ‘normalized shell density’ peaked at value = 1. Then, the GluA4: GluA2 ‘ratio of peak-normalized shell density’ was calculated for each shell of each synapse by dividing the peak-normalized blue fluorescence shell density by the peak-normalized green fluorescence shell density. The average radial distribution for each image was calculated as the mean of the individuals. Group averages for GluA3^WT^ and GluA3^KO^ across CF regions were calculated as the mean of image-means. Shells were transformed to 26 nm bins and values were plotted as a function of increasing distance from the synapse center.

For the GluA4:GluA2 fluorescence ratios in nanodomains within synapses, each synapse was divided into 9 matrices of 13×13 pixels, with eight matrices around one center matrix centered on the ribbon centroid. Thus, each nanodomain matrix was 338^2^ nm^2^, and each 9-matrix synapse (3×3 matrices) covered an area of 1,014^2^ nm^2^.

### Transmission Electron Microscopy (TEM)

A total of 8 mice (2 per genotype either in quiet or ambient) at P35 were anesthetized with a mixture of ketamine (60 mg/kg) and xylazine (6.5 mg/kg) and were transcardially perfused with 0.1 M phosphate buffer (PB) pH = 7.2, followed by the fixative containing a mixture of 3% PFA and 1.5% glutaraldehyde in 0.1 M PB. Cochleae were dissected, fixed overnight at 4°C, decalcified in 10% ethylenediaminetetraacetic acid (EDTA) for 24-48 hours at 4°C, and processed for osmication, dehydration, and embedding in epoxy resin using a similar method as previously described by Rutherford et al. (2023). A series of 15-20 ultrathin serial sections (70– 80 nm in thickness) of the mid-cochlea (20kH) were cut with a Leica EM UC7 ultramicrotome and collected on single slot gold-gilded grids with formvar. The ultrathin sections were viewed under a transmission electron microscope (TEM; JEOL Ltd., Akishima Tokyo, Japan) and IHC-ribbon synapses of modiolar and pillar side were imaged (at x40,000 magnification) with an Orius SC200 CCD camera (Gatan Inc, Warrendale, PA, USA). Only, IHC-ribbon synapses that were clearly located either on the modiolar or the pillar side of the IHC were used for further ultrastructural analysis.

### Three-Dimensional (3-D) Reconstructions and NIH Image-J Analysis of TEM micrographs

A total of 70 (GluA3^WT^ n= 35, GluA3^KO^ n= 35) IHC-ribbon synapses were used for the 3D analysis of the surface area and volume of the postsynaptic density (PSD) and the presynaptic ribbons. The TEM micrographs were calibrated, aligned, and reconstructed using the Reconstruct software (https://synapseweb.clm.utexas.edu/software-0; Fiala, 2005) as we previously described in Gómez-Nieto and Rubio (2009) and Rutherford et al. (2023). Additionally, in 27 (GluA3^KO^) and 31 (GluA3^WT^) IHC-ribbon synapses we analyzed the ribbon major and minor axes, the ribbon circularity and roundness, the PSD length, and synaptic vesicles (SVs) size from a representative TEM image of IHC-ribbon synapses using NIH ImageJ software (https://fiji.sc/) as described in Rutherford et al., (2023). SVs size was calculated as (Major diameter + Minor diameter)/2. Four to six SVs were analyzed per synapse, and the grand mean was used to compare the size of modiolar and pillar side SVs. We used the ratio of the ribbon’s major and minor axes to analyze the ribbon shape. The major/minor axes ratio denotes the shapes of the ribbons regardless of the size. Each ribbon’s ratio was classified into six groups, from 1-1.5 to >3.5. When a ribbon’s major/minor axes ratio is one, it represents a round shape, while the higher major/minor ratio represents a comparatively more elliptical shape. The presence of mitochondria in the afferent terminals of GluA3^WT^ and GluA3^KO^ in quiet was analyzed in 40 (modiolar: n= 25; pillar n= 15) and 43 (modiolar: n= 29; pillar n= 14) representative micrographs, respectively.

Representative electron micrographs of GluA3^WT^ (n= 20) and GluA3^KO^ (n= 38) in ambient were used to determine the average gray value (AGV settings: black = 0 and white = 255) per μm^2^ of afferent terminal cytoplasm, and afferent circularity and roundness with NIH ImageJ.

### Statistical Analysis

All the statistical analyses were conducted with GraphPad Prism software (version 9.4.1) or IGOR Pro (version 7.08, WaveMetrics). ABR comparisons were made with two-way mixed ANOVAs followed by Sℒidák’s multiple comparisons. Two-tailed Mann–Whitney U-test with Holm-Sℒidák’s correction for multiple comparisons or two-tailed unpaired t-tests were used to compare two independent groups. Two one-dimensional probability distributions were compared with the Kolmogorov–Smirnov (KS) test. For pairwise comparisons between more than two groups, one-way ANOVAs were used, followed by Tukey’s post-hoc correction test. The Spearman rank-order correlation coefficient was used to test the strength of the association between the two variables. Statistical significance for all the tests was set to p < 0.05. Data are represented as mean ± standard deviation (± SD).

## Supporting information

Extended view

Extended view

## Acknowledgments

This work was supported by NIDCD DC013048 (MER) and NIDCD DC14712 (MAR).

## Authors Contributions

M.E.R and M.A.R.: designed research, performed research, analyzed data, and wrote and edited the manuscript. I.P: performed research, analyzed data, wrote the first draft, and edited the manuscript. A.B.: analyzed data and edited the manuscript. M.X.: performed research and edited the manuscript. E.D.W.: analyzed data and edited the manuscript. B.V-G: designed and developed custom analytical tools.

**Data EV1.** ABR data from mice in quiet

**Data EV2**. ABR data from mice in ambient

**Data EV3**. Confocal analysis of synapses in ambient and in quiet

**Data EV4**. Confocal analysis of CtBP2m GluA2 and GluA3 at IHC-ribbon synapses in ambient and in quiet.

**Figure EV1.** Air scan confocal imaging of CtBP2 (red), GluA2 (green) and GluA4 (blue) of ribbon synapses of GluA2^WT^ (A) and GluA3^KO^ in quiet. **Support Figures 5 and 6.**

**Figure EV2.** Electron micrographs of IHC-ribbon synapses of the middle cochlear turn of female GluA3^KO^ mice in ambient at P35 (**A-C**) and P60 (**D**). **B-C**) High magnification of afferent swellings. Images A and D, are at low magnification. D’ shows high magnification of the white boxed are in D.*: afferent swellings; IHC: inner hair cell; TM: tectorial membrane. **Support Figure 7.**

