## Supplementary figures and images for "Female GluA3-KO mice show early onset hearing loss and afferent swellings in ambient sound levels"

### Extended view

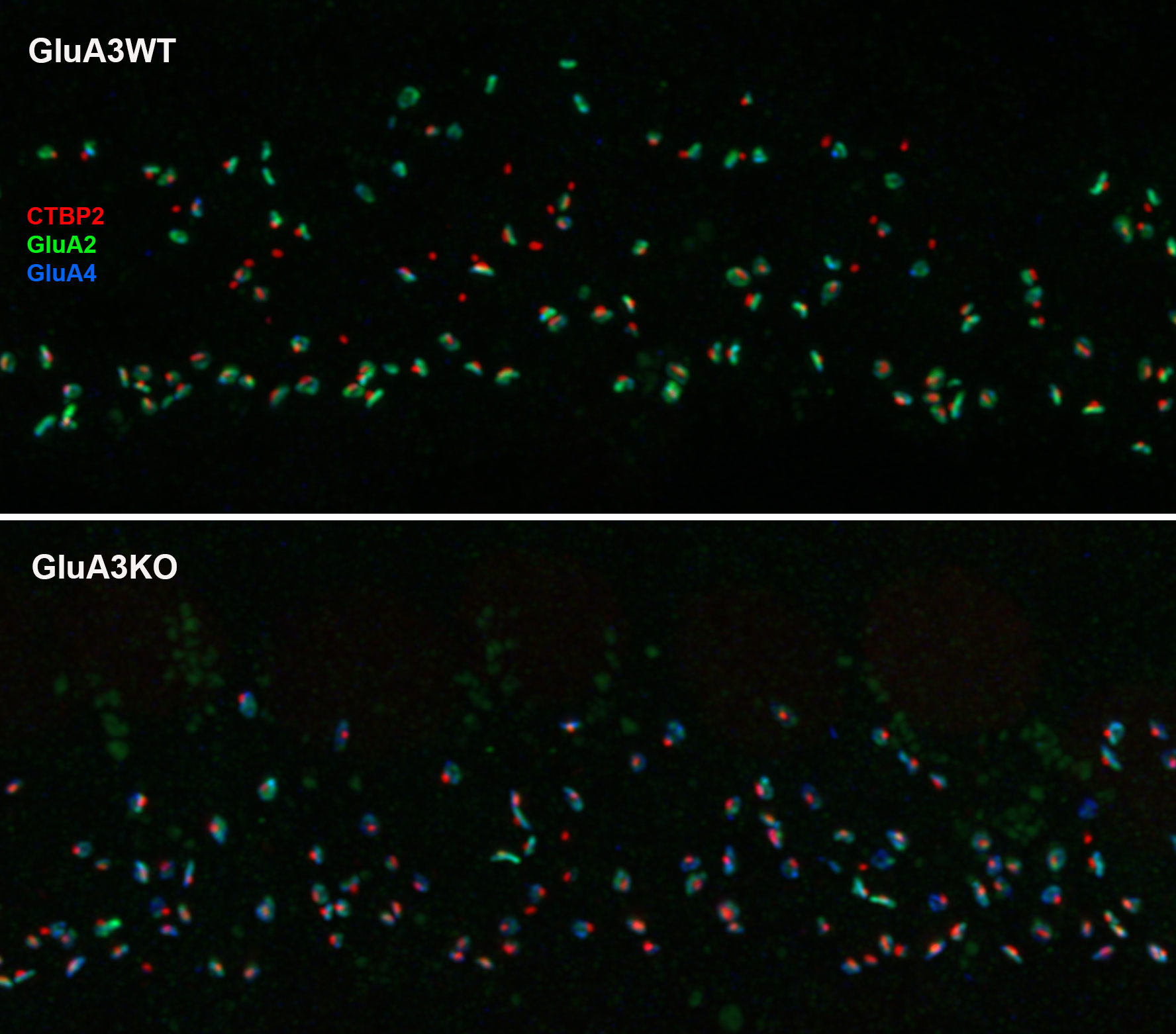

### Extended view

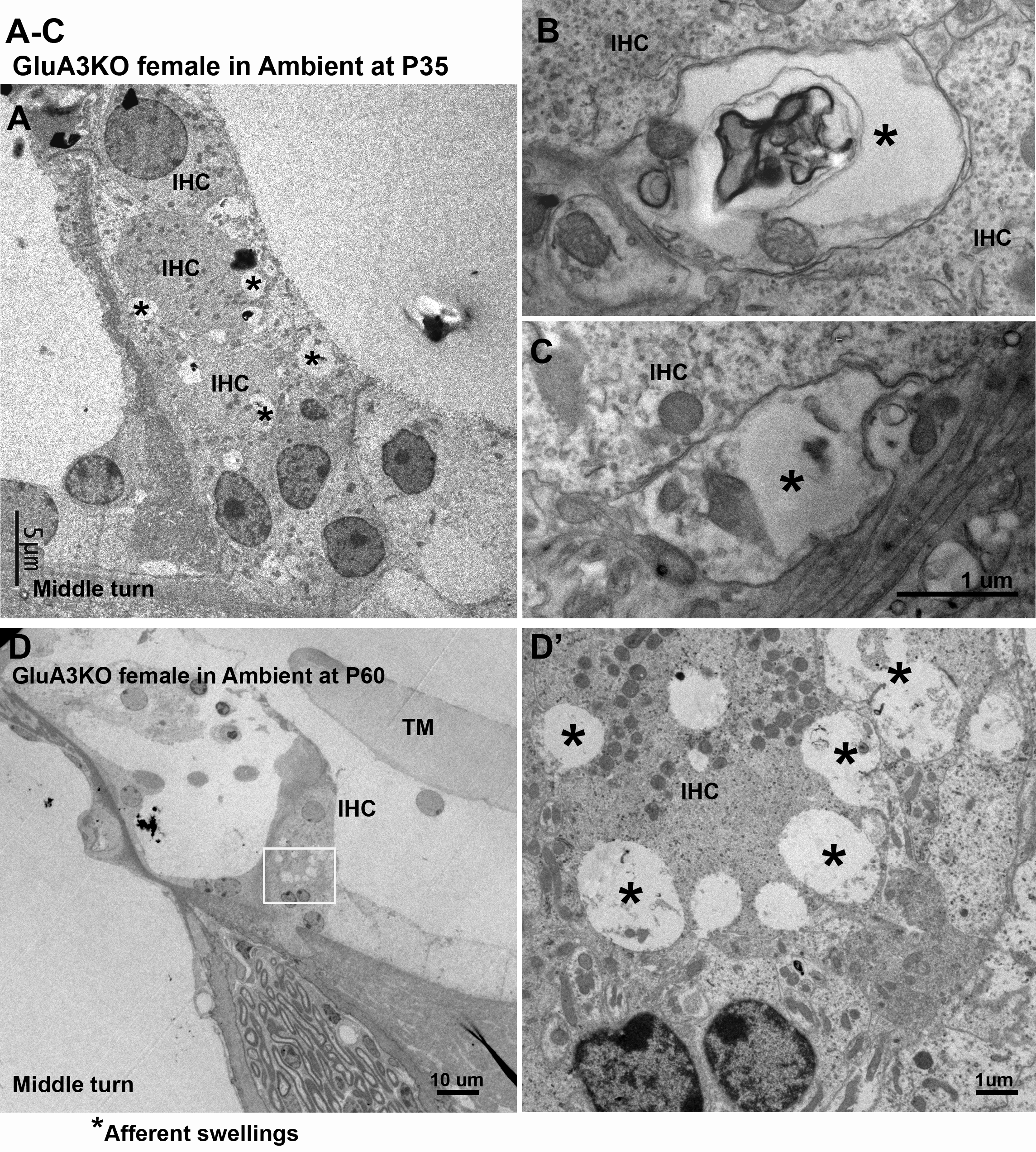
